# TooManyCells identifies and visualizes relationships of single-cell clades

**DOI:** 10.1101/519660

**Authors:** Gregory W. Schwartz, Jelena Petrovic, Maria Fasolino, Yeqiao Zhou, Stanley Cai, Lanwei Xu, Warren Pear, Golnaz Vahedi, Robert B. Faryabi

## Abstract

Transcriptional programs contribute to phenotypic and functional cell states. While elucidation of cell state heterogeneity and its role in biology and pathobiology has been advanced by studying single cell level measurements, the underlying assumptions of current analytical methods limit the identification and exploration of cell clades. Unlike other methods, which produce a single uni-layer partition of cells ignoring echelons of cell states, we present `TooManyCells`, a software consisting of a suite of graph-based tools for efficient, global, and unbiased identification and visualization of cell clades while maintaining and presenting the relationship between cell states. `TooManyCells` provides a set of tools based on a matrix-free efficient divisive hierarchical spectral clustering algorithm wholly different from the prevalent Louvain-based methods. `BirchBeer`, the visualization component of `TooManyCells`, introduces a new approach for single cell analysis that is built on a concept intentionally orthogonal to the widely used dimensionality reduction methods. Together, this suite of tools provide a paradigm shift in the analysis and interpretation of single cell data by enabling simultaneous comparisons of cell states at context-and application-dependent scales. A byproduct of this shift is the immediate detection and visualization of rare populations that outperforms previous algorithms as demonstrated by applying these tools to existing single cell RNA-seq data sets from various mouse organs.

## Introduction

Cellular transcriptional output contributes to cell type and state determination. Emergent technologies such as single cell RNA-seq (scRNA-seq) have improved identification and characterization of cell state heterogeneity driven by differences in gene expression programs and have advanced our understanding of the functional role of cell diversity in domains such as cell differentiation, response to stimuli, and dysregulation in diseases.

Many clustering algorithms propose partitioning of scRNA-seq data in an effort to identify groups of cells with related transcriptional programs, fractionate cells into known cell types or states, and in some cases delineate novel rare cell populations. Louvain-based clustering methods^1^ are among the most widely used scRNA-seq clustering approaches. These algorithms iteratively optimize a measure of community, Newman-Girvan modularity, in a greedy manner to generate a single partitioning of cells. In most scRNA-seq analysis workflows, the identified cell clusters with similar transcriptomic patterns are then visualized using dimensionality reduction techniques such as t-distributed stochastic neighbor embedding (t-SNE)^2^, where a high dimension gene expression space is stochastically reduced to project the locations of each individual cell onto a two dimensional surface. This workflow leads to the identification of a single uni-layer partition of cells that are interpreted using visualization methods lacking quantitative inter-and intra-group relationship information. Given that precise cell state definitions and their inter-and intra-relationships are context-dependent and might not fit into a single uni-layer partition of cells but rather a hierarchical structure of nested cell states, we introduced a paradigm shifting suite of tools for the analysis, interpretation, and visualization of single cell data by enabling simultaneous delineation and visualization of multi-layer cell clades.

To elucidate cell state diversity and enable interpretation of inter-and intra-state relationships, we proposed an entirely different approach for single cell analysis that was specifically developed to handle flexible and multi-scale cluster definition and interpretable visualization. To this end, we developed `TooManyCells`, a suite of graph-based algorithms and tools for efficient, global, and unbiased identification and visualization of cell clades while maintaining and presenting cell groups’ interrelationships. `TooManyCells` provides a novel single cell clustering algorithm `ClusterTree` that implements an efficient matrix-free divisive hierarchical spectral clustering for single cell measurement analysis. In addition, we implemented methods in `TooManyCells` that enable the assessment of cell diversity and estimate the sample size required to survey a specified level of cell species richness in a heterogenous population.

`TooManyCells` also provides `BirchBeer` which is a fully customizable novel model for visualization of hierarchical cell clustering analyses at single-cell resolution. While rendering the clustering tree, `BirchBeer` makes use of weighted average blending of coloring labels, scaling branches, modularity overlays, internal node labeling, leaf node summarization, and more to enable multilayer and multifaceted exploration of single cell clades. To provide maximum applicability, `TooManyCells` loads and generates several standard file formats. Consequently, alternative clustering algorithms can be used in tandem with the visualization, providing a harmony between multiple approaches to assist in elucidating single cell relationships. Together, the proposed clustering and visualization algorithms enable delineation of context-and application-dependent cell clusters and expedite the exploration and quantitation of interrelationships among cell clades.

We demonstrated the effectiveness of `TooManyCells` in reliably identifying and clearly visualizing groups of abundant and rare populations using several analyses and benchmarks. Publicly available single cell data sets from 11 mouse organs as well as controlled subsetting and mixing experiments of single cell populations were used to benchmark `TooManyCells`’ performance against several widely used single cell clustering algorithms. Our data showed that `TooManyCells` recapitulates known relationships between cell types within mouse organs. Interestingly, where other popular methods fail, `TooManyCells` detects two rare populations each down to the smallest benchmark of 0.5% prevalence in the cell admixture. Additionally, `TooManyCells` assisted in a fine grain B cell subset definition within mouse spleen scRNA-seq data set, where a plasmablast population was overlooked using a popular Louvain-based algorithm. `TooManyCells` and its individual components are available through https://github.com/faryabiLab/too-many-cells.

## Results

### TooManyCells for clustering and visualization at single-cell resolution

`TooManyCells` is a suite of single cell analysis tools for identification and visualization of cell clades (Figure 1a). `TooManyCells` provides novel algorithms for clustering, visualization, clumpiness, diversity quantification, and rarefaction analyses of single cell measurements. In addition, a number of auxiliary functions are integrated in `TooManyCells` for flexible data normalization, differential expression analysis, and data import and export to name a few.

**Figure 1:**
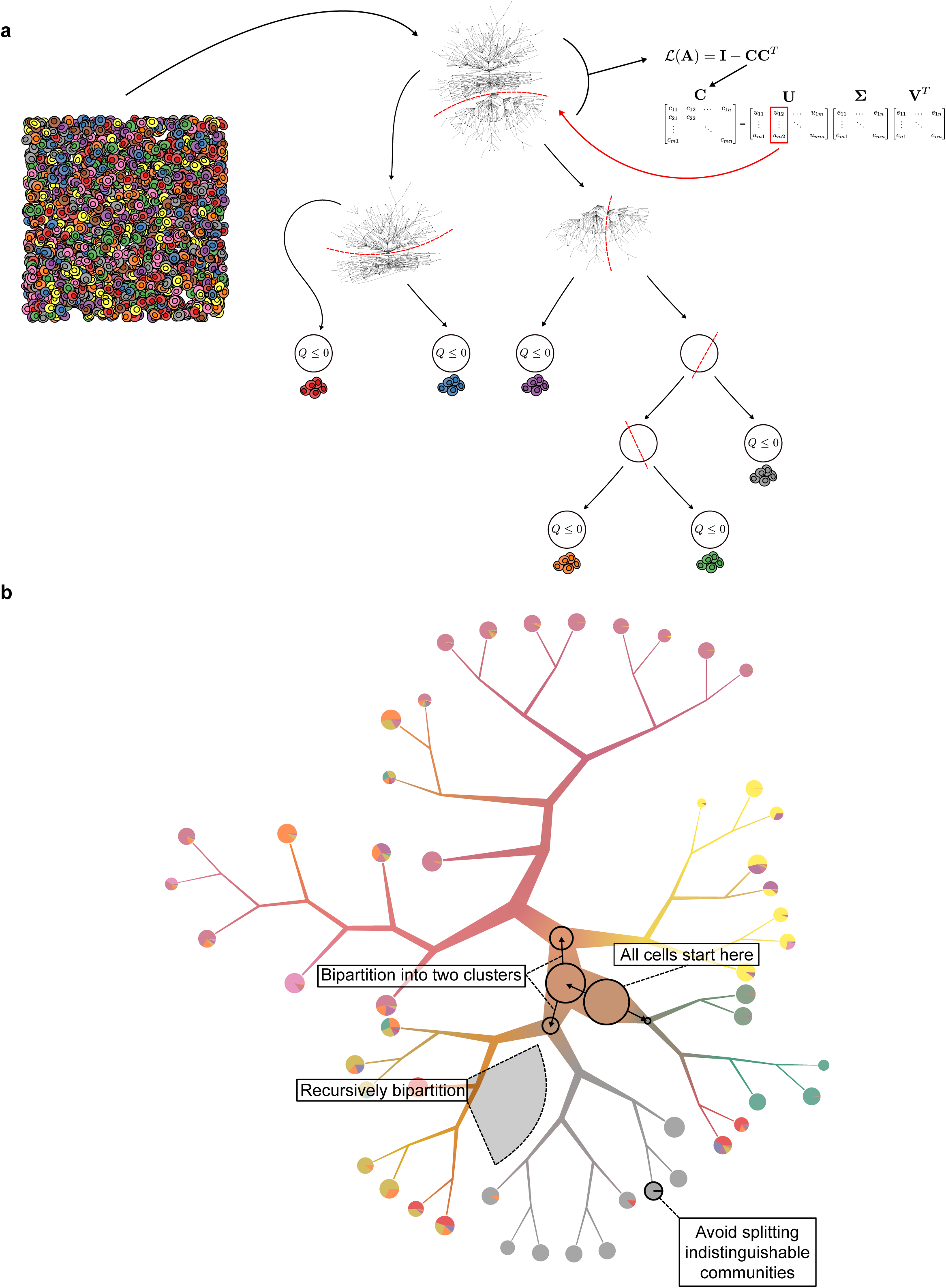
`TooManyCells` process. (a) Divisive hierarchical spectral clustering. The network of nodes (cells) and edges (cosine similarity) is recursively bi-partitioned using a matrix-free with truncated-SVD calculation of the singular vector of the second largest singular value of a new matrix **C** until a candidate split results in non-positive Newman-Girvan modularity (*Q*). (b) `BirchBeer`, a visualization component of `TooManyCells`, visualizes divisive hierarchical spectral clustering of single cells while providing many capabilities and options including but not limited to weighted average blending of colors, scaling branches, modularity overlays, smart tree pruning, leaves node visualization options.

While `TooManyCells` enables an end-to-end builtin scRNA-seq analysis solution, it also provides a degree of modularity for more flexible usage. Many of the provided options are built into the program and can be disabled completely at the command line interface to allow for other preprocessing procedures, including initial filtering, normalization, similarity measure calculation, graph construction, clustering, and more. For instance, to enable an end user to analyze any graph, such as a nearest neighbor graph with a different similarity measure, `TooManyCells` provides a standalone implementation of its clustering algorithm `ClusterTree` that computes on any similarity matrix. Data visualization at single-cell resolution is an integral part of scRNA-seq data interpretation. To this end, a fully customizable novel model for visualization of hierarchical cell clustering analyses at single-cell resolution, called `BirchBeer`, was implemented in `TooManyCells`. `BirchBeer` is designed to display any tree data structure independent of the clustering algorithm to enhance the versatility of data visualization. Many components of `TooManyCells` are developed in independent libraries that can be altered or incorporated into other scRNA-seq analysis solutions. All software is available for installation as libraries, programs, and Docker images for `TooManyCells` and `BirchBeer`.

### ClusterTree: a matrix-free divisive hierarchical spectral clustering algorithm for scRNA-seq

As cell states are regulated by continuous gene expression levels, we reason that the separation of cells into groups with related expression programs would be more accurately represented using a mathematical framework that provides interrelated multi-layer cell clades without any predefined cell population. For instance, recursive partitioning of a hierarchical structure from scRNA-seq measurement can recapitulate lymphoid populations in spleen where a bipartition separates B from T cells, then plasma cell lineage could be separated from transitional B cells followed by factionating of plasmablasts and plasma cells till reaching the separation of plasma cells with different B cell receptor structures. To identify a continuum of cell states, `ClusterTree`, the clustering component of `TooManyCells`, starts with all cells belonging to single group acting as the root of the clustering tree (Figures 1a and 1b). These cells are represented by an *m* × *n* matrix with *m* cells and *n* genes. Each entry in the matrix is the abundance data for a gene in a cell.

`ClusterTree` uses spectral clustering to partition the cells into two groups. Traditionally, spectral clustering with normalized cuts uses the eigenvector corresponding to the second smallest eigenvalue of the normalized Laplacian matrix (See Methods). However, the normalized Laplacian matrix requires the calculation of *m* × *m* adjacency matrix where each entry denotes the similarity between two cells. Obtaining and factorization of adjacency matrix impose significant time and memory requirements on each iteration of recursive spectral clustering^3^. To mitigate this limitation, a matrix-free spectral clustering algorithm is implement in `ClusterTree`. This algorithm instead calculates the singular vector of the second largest singular value using truncated singular vector decomposition (SVD) of a new matrix, **C** to bipartition the cells^3^ (See Methods).

Briefly, at each recursive step, `ClusterTree` efficiently identifies the candidate cell bipartitions and evaluates its significance using Newman-Girvain modularity (*Q*)^4^. `ClusterTree` recursively bipartitions cells until the separation of cells into two groups is as significant as creation of two cell communities randomly, i.e. *Q* < 0. At this condition, the cells form a fine-grain collection as a leaf node of the clustering tree. Use of the modularity as a stopping criteria instead of an optimization parameter bypasses limitations associated with global optimization-based clustering^5^. As such, `ClusterTree` produces a hierarchy of nested cell clusters where each inner node is a cluster at a given scale and a leaf node is a fine-grain cluster where any additional bipartitioning would be as good as a random splitting of the cells (Figure 1).

### BirchBeer: visualization model for scRNA-seq data interpretation

Visualization at single-cell resolution is an integral part of scRNA-seq data interpretation. To accommodate for interrogation of a hierarchy of nested cell clusters, `TooManyCells` is equipped with `BirchBeer`, a fully customizable novel software developed specifically for clear and interpretable display of hierarchical structures of cell clusters (Figure 2a). `BirchBeer` is specially designed to display the entire tree with several additional features that assist the users with finding relevant populations and facilitating data exploration. `BirchBeer` can scale each branch according to the number of cells in that subtree and color each branching point using the weighted average blend of cell label colors connected through that branch (Figure 2a). The color blending could enhance the identification of overlapping or distinct populations throughout multiple scales (Figure 2b). Multiple visualization formats are available for a cell group at leaf nodes. A leaf node can present a pie chart showing the distribution of its cell composition. To depict information at the single-cell resolution, each individual cell can be rendered and color-coded at a leaf node to visualize additional information such as the exact number of cells, the expression level of a given gene at each cell and more in a leaf cluster (Figure 2b).

**Figure 2:**
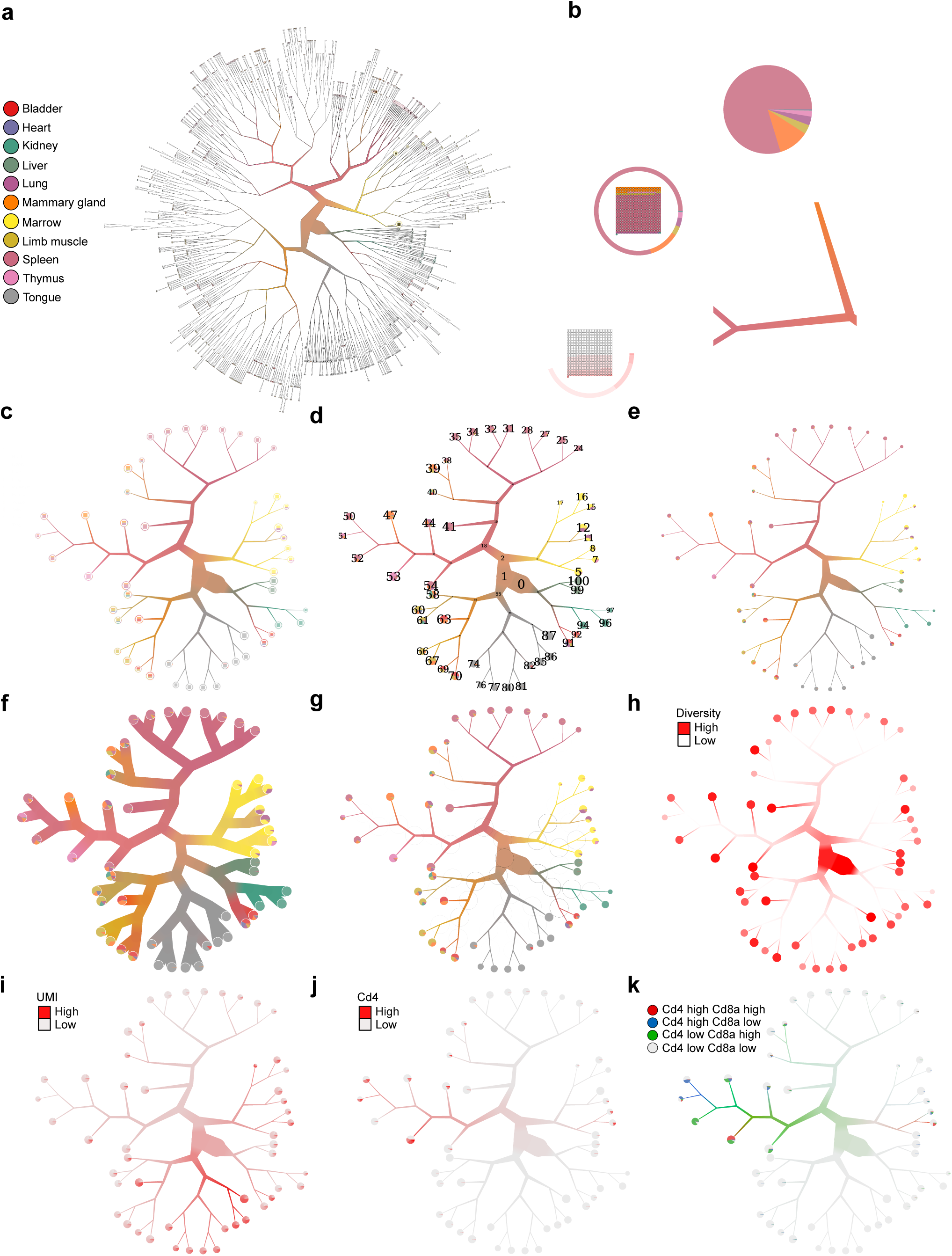
Example of `BirchBeer` visualization capabilities. (a) The complete tree with default settings. (b) Different leaf rendering options (clockwise from bottom: gene expression, “pie ring”, pie chart) and example of scaling and average weighted color blending for branches. (c) Tree from (a) with cutting criterion `--smart-cutoff 4`, which is used in all the following panels d-k. (d) Tree with numbered nodes. (e) Tree with non-default scaling width (f) Tree with disabled branch scaling. (g) Tree with modularity of bi-partitioning at each internal node displayed as black circles, where higher modularity is represented by darker circumference intensity. (h) Color-coded tree with a continuous variable (e.g. cell diversity of organs, Increasing color intensity represents increasing diversity). Inner and leaf nodes use different intensity scales. (i) Color-coded tree with a discrete variable presenting UMI counts. (j) Color-coded tree with expression level of a specific gene (e.g. *Cd4* expression level). (k) Color-coded tree with expression level of multiple genes (e.g. *Cd4* and *Cd8a* expression level).

For larger and complex cell admixture, t-SNE and similar projection methods suffer from the rendering of many overlapping cells that overwhelm the single-cell resolution visualization. The lower branches and leaf nodes of `BirchBeer` clustering tree may also be visually obscure due to the plethora of leaf nodes (Figure 2a). Given that `TooManyCells` divisive hierarchical clustering algorithm produces a nested cluster structure where relationships between the groups are maintained, the inner nodes also represent cell clusters at coarser scales. To assist with interpretation and analysis of complex cell admixture, `birch-tree` provides several methods for pruning clustering tree (Figures 2c-k). These methods include the number of steps away from the root, the minimum leaf node size, and a quantile-based pruning solution (Figure 2c). Importantly, `TooManyCells` enables comparison between both the leaf node clusters and all the nested clusters in the hierarchy of clusters. To assist in selecting a given cell partition, `BirchBeer` provides an overlay of each inner or leaf node cluster identification number that can be used as an input to the `TooManyCells` differential expression analysis module for comparative analysis of clusters at different scales or other downstream analyses (Figure 2d). As branch scaling widths may be aesthetically subjective, they can be adjusted (Figure 2e versus Figure 2c) or entirely disabled (Figure 2f). As the modularity of a candidate bipartitioning demonstrates the significance of that split relative to the random division model, `BirchBeer` can display the modularity as black circles of varying degree of circumference darkness to aid in identifying populations that are very distinct from each other (Figure 2g).

Overlaying continuous or discrete readouts could be used to further explore clusters identified in a scRNA-seq analysis. For instance, a quantitative measure of cell admixture diversity can be used to quantify the number of cell subpopulation in a given cluster. To assist in finding tree locations of high or low diversity, `BirchBeer` can visualize diversity of inner and leaf nodes (Figure 2h). In addition to overlaying continuous variables on a tree, `BirchBeer` allows for overlaying discrete variables. For instance, `BirchBeer` can overlay the UMI counts for each cell and cluster (Figure 2i), as well as the expression status of one or more genes to accommodate for visual cell fractionation analyses (Figures 2j and 2k).

### TooManyCells identifies pure cell clusters

To assess the performance of `TooManyCells`, we used the *Tabula Muris* data sets^6^ to examine the extent of cell homogeneity in clusters identified by `TooManyCells` and other popular clustering algorithms. As part of the *Tabula Muris*, 11 organs were profiled by scRNA-seq in three month old mice. Assuming that the cell clusters should mainly comprised of cells from an organ, cluster purity was assessed using order 1 diversity index^7–9^, which quantifies the effective number of organs represented in a cluster (see Methods). We compared the diversity of clusters identified by `TooManyCells` to those of Cellranger, Monocle^10^, Phenograph^11^, and Seurat^12^ (Figure 3). The default filters and parameters of each algorithm were considered. Given that `TooManyCells` produces a hierarchy of clusters and not just a single partition of cells as the other methods do, cutoffs at several levels of the clustering tree were considered to have a balanced comparison. A comparative analysis was performed based on an increased level of cell mixture complexity, where the first 3, the first 6, the first 9, and finally all 11 data sets from thymus, spleen, bone marrow, limb muscle, tongue, heart, lung, mammary gland, bladder, kidney, and liver were considered.

**Figure 3:**
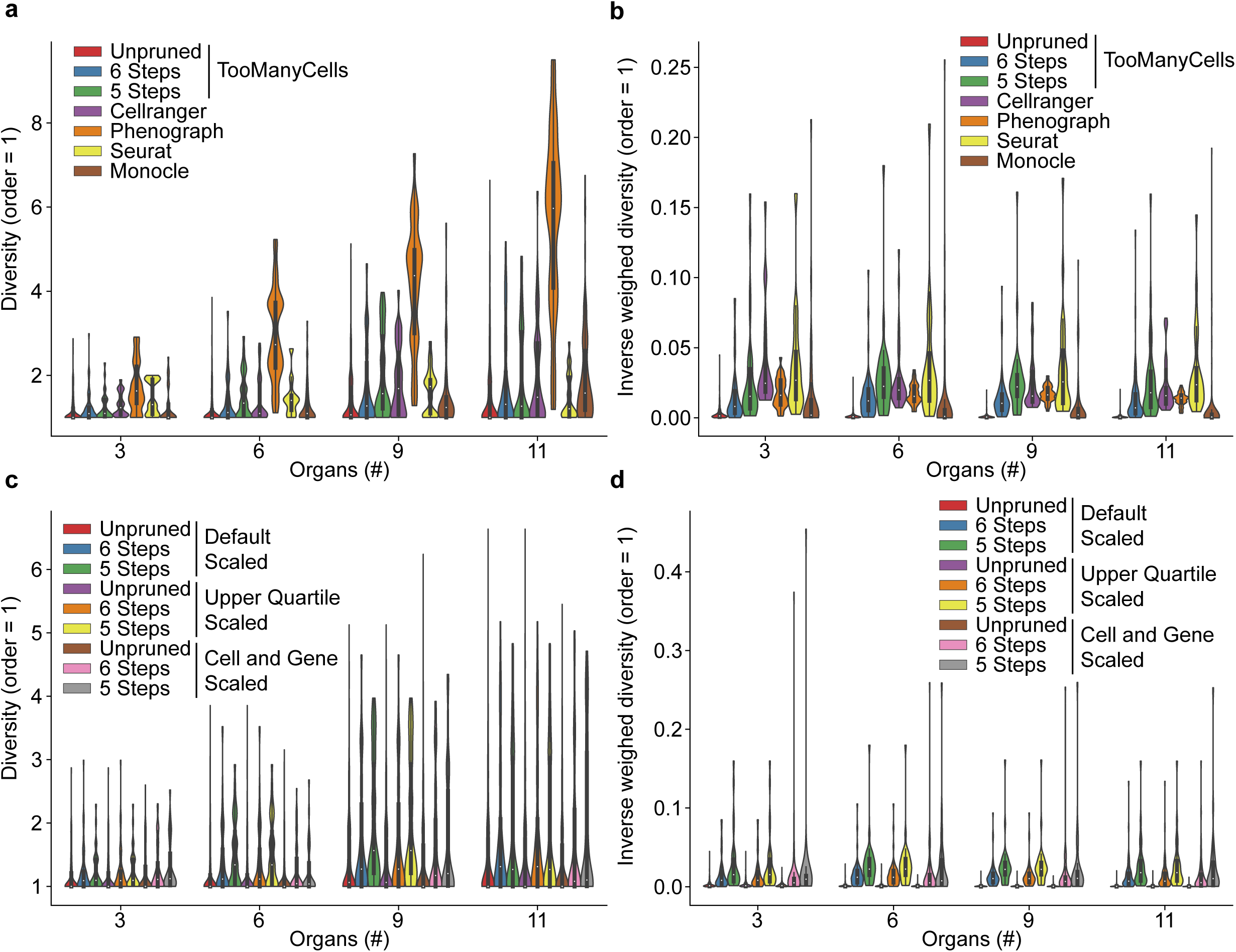
Comparative analysis of cluster purity. (a) The population analyzed is represented by the number of sites (i.e. organs and tissues) on the x axis taken from the left of the ordering {Thymus, Spleen, Marrow, Limb muscle, Tongue, Heart, Lung, Mammary gland, Bladder, Kidney, Liver}. The distribution of cluster diversity was compared. Either all, 64, or 32 (all steps from root, 6 steps from root, or 5 steps from root) clusters from the `TooManyCells` hierarchical cluster tree were considered. Default filterings and parameters were used for all algorithms. Higher diversity represents lower accuracy. (b) The inverse weighted diversity is defined as the product of inverse min-max normalized diversity and the ratio of cluster size to the total number of cells. Larger values indicate purer and larger clusters. (c) Same as (a) for different `TooManyCells` normalization procedures: the `TooManyCells` default normalization (Default Scaled) term frequency-inverse document frequency (tf-idf), upper quartile normalization (Upper Quartile Scaled) and total and gene median count normalization (Cell and Gene Scaled) followed by tf-idf normalization. (d) Same as (b) for different `TooManyCells` normalization procedures.

In all four data complexity levels, Seurat resulted in more pure clusters with a lower median diversity compared to Phenograph, even though both use Louvain clustering to optimize Newman-Girvain modularity (Figure 3a). Cellranger’s algorithm outperformed Phenograph and Seurat for the mixture of the first 3 and 6 organ data sets but was inferior to Seurat when the complexity of data increased to 9 or all 11 organs (Figure 3a). Monocle produced a large number of purer small clusters for low complexity experiments (3 and 6 sites in Figure 3a), while exhibiting increased diversity over Seurat for higher number of organs admixture data. Together, these results suggest that Monocle outperforms Cellranger, Phenograph, and Seurat when data sets are less complex and more homogeneous, while performing worse than Seurat for more heterogeneous and diverse data sets.

Our results showed that when no clustering depth limitation was imposed and `TooManyCells` partitioned the cells into more refined clusters, the algorithm always was one of the best performing algorithms to delineate pure clusters. Yet, `TooManyCells` produced more diverse clusters consisting of cell mixtures from more organs when the clustering depth was restricted (depth of 5 or 6 from the root of the hierarchical tree, i.e. TooManyCells 5 and 6 steps, respectively). Limiting the `TooManyCells` clustering steps from the root forced the algorithm to prematurely terminate the cell partitioning. We observed that `TooManyCells` with less clustering depth exhibited median cluster purity comparable with other methods for the less complex cell mixtures, while performing only better than and on par with Monocle and Phenograph, respectively, for the complex 11 organs data mixture (Figure 3a TooManyCells 5 and 6 steps). As such, the `TooManyCells` flexibility and multi-layer clustering output enables context-dependent clustering results from the entire hierarchy of the presented clusters according to a desirable purity level. The choice of clustering granularity could be guided by the visualization from `BirchBeer` that could contextualize cluster features such as relative size, modularity (Figure 2h), and distance from the root.

While diversity index quantifies the ability of clustering algorithm to produce groups of cells with similar expression patterns, this measure does not account for differential cluster sizes produced by various methods. As such, a clustering algorithm that produces arbitrarily small clusters could potentially partition homogenous cells into unnecessarily smaller groups with very high purity as opposed to outputting similarly homogeneous cluster of a larger size with similar cell homogeneity. To account for both the cluster size and diversity, we compared clustering algorithms based on the inverse weighted diversity defined as the product of inverse min-max normalized diversity and the ratio of cluster size to the total number of cells, with better performance represented by larger values (Figure 3b). The two algorithms that could generate smaller size clusters, Monocle and `TooManyCells` without any clustering depth limit, resulted in lower inverse weighted diversity. While, `TooManyCells` with limited clustering steps rivaled Seurat, which tends to produce larger size clusters (Figure 3b). Together, these data showed that in contrary to other algorithms such as Monocle and Seurat that produce either small or large size clusters, multi-layer nested `TooManyCells` clusters provide flexibility to control clustering granularity and enable intuitive trade-offs between cluster purity and size.

We further considered different possible normalizations to evaluate the robustness of `TooManyCells` to data normalization with respect to both cluster purity and size. To this end, three normalization approaches were considered: 1) term frequency-inverse document frequency (tf-idf)^13,14^ normalization alone, 2) cell normalization by total count and gene normalization by median count followed by tf-idf, 3) tf-idf with preceding upper quartile normalization (See Methods). In general, `TooManyCells` performance was just marginally influenced by the choice of data normalization (Figures 3c and 3d). As expected, upper quartile normalization had no effect on the `TooManyCells` clustering performance as cosine similarity is robust to multiplicative scalings (Figures 3c and 3d). Using total count and gene median count normalization resulted in a slight performance improvement for `TooManyCells` with a limited clustering depth as measured by both the cluster diversity and inverse weighted diversity (Figure 3c, and 3d). Yet, this normalization had an insignificant effect on `TooManyCells` performance without clustering depth limitation. Together, these comparative analyses suggest the importance of assessing clustering algorithm performance and motivates a careful choice of clustering methods according to both cluster purity and size metrics. `TooManyCells` visualization model `BirchBeer` enables rendering these metrics to improve the interpretability of scRNA-seq clustering results.

### TooManyCells accurately delineates rare populations in cell admixtures

The detection of rare cell populations is one of the major motivation of single cell transcriptomic analysis. While many single cell clustering algorithms claim to identify rare populations, few have explicitly benchmarked this ability. In order to test the affinity of algorithms for finding rare populations in a rigorous and robust manner, we simulated different levels of rare and common populations based on the actual biological data from the mouse organs. An accurate single-cell resolution clustering is expected to not only detect the rare populations from the common but also distinguish the rare populations from each other. To this end, two rare populations of equal sizes were mixed with a common cell population. Specifically, three set of cells from the mouse tongue, bone marrow, and mammary gland were considered. Cells from tongue were used as common cells and mixed with rare populations from bone marrow and mammary gland. Tongue cells were dissimilar between both marrow and mammary gland cells (Figure S22), encouraging the latter samples to behave as outlier rare populations. In ten different simulation settings a total of 1000 cells were mixed ranging from 900 to 990 common cells and 100 to 10 rare cells (half from bone marrow and half from mammary gland). For example, in the first setting there were 1000 cells comprised of 90% common and 10% rare (5% of each rare population), while in the tenth setting the 1000 cells comprised of 99% common and 1% rare (0.5% of each rare population). Visual inspection of t-SNE projections showed apparent discrepancies between the actual origin of cells and their cluster labels (Figures 4a and 4b). Regardless of the clustering algorithm, the t-SNE plots were limited in distinguishing the two rare populations in a cell admixture. Upon initial visual inspection of the t-SNE plots, one could speculated that any of the small cell islands could contain the rare populations (Figures 4a and 4b left columns). However, there are several notable islands in these plots and, without prior labeling of the organs, it would be difficult to distinctly identify the rare mammary gland and bone marrow cells. This issue is inherent to a t-SNE plot, where both distance and density are converted to local density.

**Figure 4:**
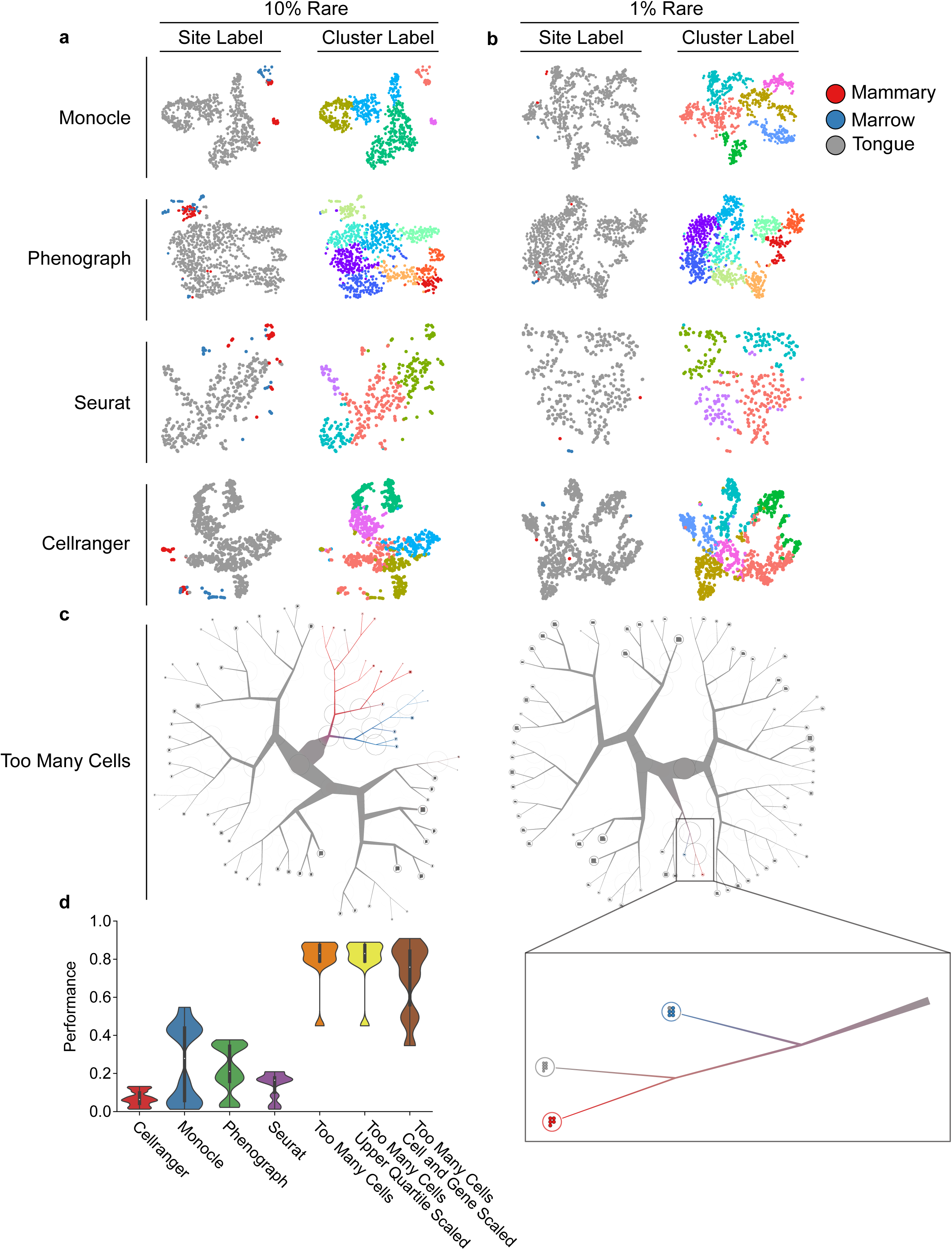
Detection of cells from two rare populations mixed with a common cell population was benchmarked for common clustering algorithms (a and b). Columns from left to right: cells labeled by their actual types (i.e. organs) and cells labeled by their assigned clusters of a scRNA-seq clustering algorithm. Rows from the top to bottom: Monocle, Phenograph, Seurat, and Cellranger t-SNE projections. The analysis for a) 900 common (tongue) and 100 rare (50 mammary gland and 50 marrow) cells and b) 990 common and 10 rare cells (5 mammary gland and 5 marrow) are presented. (c) The result of `TooManyCells` clustering and visualization. Left: 900 common (tongue) and 100 rare. Right: 990 common and 10 rare cells. (d) Quantification of rare population benchmark. For each run, the performance represents true rare pairs (mammary gland with mammary gland and marrow with marrow in the same cluster) / total rare pairs (true rare pairs and mammary gland with marrow).

By contrast, the `TooManyCells` visualization component `BirchBeer` is specifically designed to plot cluster relationships and thus readily presented the rare populations at single-cell resolution. For 90% to 10% mixing of the common and rare populations, `TooManyCells` separated the rare and common populations in the first tree bifurcation, followed by spitting the two rare groups in the next branching of the subtree (Figure4c left panel). Interestingly, rare populations would still be easily identifiable in the absence of their labels as they were separated in the first branching from the main node which was quite small in comparison to the other subtrees, as visualized by the branch thickness (Figure4c left panel). With modularity overlaying the nodes, the rare populations were even more apparent as the maximum modularity of separation appeared at these locations. For 99% to 1% mixing of the common and rare populations, `TooManyCells` again delineated the rare populations relatively close to the root of the clustering hierarchy and readily presented the cells of the rare populations with the help of a drastically smaller subtree compared to the other side of the branching (Figure 4c right panel).

To quantitatively compare the performance of `TooManyCells` and other commonly used clustering algorithms in detecting rare populations, a contingency table of the fraction of pairwise labels was generated from the clustering result of each mixing population. A pairing of two cells from different sites (e.g. mammary gland cell with marrow cell) was counted as a false assignment, while a pairing of two cells from a same site was classified as a true one (e.g. a mammary gland cell with another mammary gland cells or a marrow cell with another marrow cell). The fraction of true pairs in all pairs was used as a performance measure. Each mixing population was analyzed with all the algorithms studied in the cluster purity assessment (Figure 4d, Cellranger: Figure S1, Monocle: Figure S2, Phenograph: Figure S3, Seurat: Figure S4, `TooManyCells` with various normalization methods: Figure S5 and Figure S6). Quantifying these results showed that although Seurat performed well according to cluster purity metric, this algorithm performed poorly in detecting rare populations, second to Cellranger for the worst accuracy (Figure 4d). Similarly, Phenograph exhibited the third worst performance, perhaps due to the usage of Louvain clusign in both algorithms. While Monocle detected rare populations more accurately than Phenograph and Seurat, `TooManyCells` with any normalization method significantly outperformed all algorithms, with the sole tf-idf normalization and tf-idf with preceding upper quartile normalization being the most performant (Figure 4d).

### TooManyCells elucidates relationships between mouse cell types

The complexity of cell composition varies across tissues and organs. To explore how known biologically distinct cell types related to the identified cell clusters defined by our scRNA-seq analysis, we applied `TooManyCells` to each of the 11 mouse organs separately (Figures S7 – S17). Using cell annotations from The Tabula Muris Consortium^6^, we observed similar cell types aggregating together within the trees, in concordance with the t-SNE projections from the Tabula Muris (Figures S7 – S17). Using these trees, we next compared the heterogeneity of cell clusters at the most refined level of `TooManyCells`-generated hierarchical outputs with the previously annotated cell populations. To this end, we used the diversity index to quantify and compare the heterogeneity of the most refined level of `TooManyCells`-generated hierarchical outputs with cell diversity labeled by the cell annotations defined by known cell markers^6^ (Figure S18a, See Methods). We observed a significantly lower diversity using these cell markers, suggesting that current known cell markers potentially lack the resolution to capture the underlying cellular diversity and the cell state continuum comprising each of these mouse organs.

Observing a disparity in cell heterogeneity of various mouse organs instigated us to equip `TooManyCells` with an algorithm that enables informed and cost-effective scRNA-seq experimental design. To this end, `TooManyCells` provides rarefaction curve analysis to estimate and validate sufficient sample size required to survey cell diversity given the level of admixture heterogeneity (Figure S18b, see Methods). Refraction analyses of 11 mouse organs showed that the Tabula Muris data sets provide sufficient single cell measurements to fully saturate detection of diverse cellular states in each of these organs, and profiling 3000 cells was sufficient to capture the heterogeneity of these organs using `TooManyCells` clustering algorithm. Rarefaction analysis using the known cell markers as labels demonstrated a decreased sampling requirement to fully saturate observation of cell type (Data not shown). These result imply that less cells are needed to observe every *a priori* annotated cell type than cell state defined by unsupervised exploration of cell state continuum.

Despite a high degree of complexity, the 11 mouse organs share potentially similar cell types. To investigate the overall similarities between the cellular composition of these organs as measured by scRNA-seq, we used a “clumpiness” measure, quantifying the degree of aggregation of labels within a hierarchical clustering structure^15^ (see Methods). To this end, if cells from an organ are uniformly dispersed throughout the entire tree, that organ would have a low clumpiness. Conversely, if all the organ cells aggregate within a subtrees then that organ would be characterized by a high degree of clumpiness. We used the clumpiness measure after labeling the cells in the `TooManyCells`-generated hierarchical clustering by either their site of origin (Figure 2a) or previously annotated cell types within each organ (Figure S19 and Figure S20). We first used the clumpiness measure with cell organs as labels (Figure S21) in order to quantify the relationship between the cell state composition of different organs with respect to the `TooManyCells`-generated hierarchical cell clusters (Figure 2a). The clumpiness of organ labels showed that cells from thymus, spleen, and bone marrow aggregate together within the hierarchical cell clades of the 11 mouse organs (Figure S21), possibly due to their enrichment for of various hematopoietic cell types. Furthermore, the cells from the mammary gland and limb muscle were the most closely related cells, which could be due to the presence of fibrous connective tissues in these organs. Finally, cell composition of tongue and liver were the most dissimilar to all the other 11 mouse organs (Figure S21).

We next investigated whether `TooManyCells`-generated hierarchical cluster tree could capture the relatedness of similar immune cell types located in different organs (Figure S22). The clumpiness measure showed that both T (Figure 2j) and B (Figure S23) cells from various organs are related with the notable exception of developing lymphocytes such as the pre-pro B cells and B cells from the bone marrow, which were found together in another section of the `TooManyCells`-generated hierarchical cluster tree (Figure S22). Interestingly, various immune cells from the lung were grouped together and were separated from their counterparts in other organs (Figure S22). With the exception of lung and bone marrow residing macrophages, the clumpiness analysis showed that macrophages were strongly related across various organs (Figure S22). Together, these data demonstrate that `TooManyCells` not only identifies and intuitively visualizes the hierarchy of cell state clades from scRNA-seq measurements, but also provides clumpiness and diversity measures to interrogate relationships and properties of cell heterogeneity from a clustering tree respectively.

### TooManyCells identifies plasmablasts in mouse spleen

Plasmablasts are a rare B cell population in the spleen^16^. To further demonstrate the ability of `TooManyCells` to stratify rare groups of cells, we analyzed the immune cell composition of the mouse spleen. This analysis showed that `TooManyCells` readily separated B cells, T cells, macrophages, and dendritic cell populations Figure 5a). Importantly, B cells and T cells comprised the majority of the data set and were represented in the tree at the first initial branch point. The macrophages were less abundant population that were separated from the T cells and were further divided into more subgroups in the tree with high degree of modularity, suggesting high heterogeneity of these cells. Similarly, dendritic cells were also partitioned in high modularity locations throughout the tree (Figure 5a). We next used prior RNA biomarkers and B cell biomarkers as defined by ImmGen^17^ to further identify B cell subtypes (Figure 5b). Interestingly, this refined labeling of the `TooManyCells` generated tree of mouse spleen led to the identification of a branch enriched for a plasmablast signature versus other B cell subtypes (Figure 5c).

**Figure 5:**
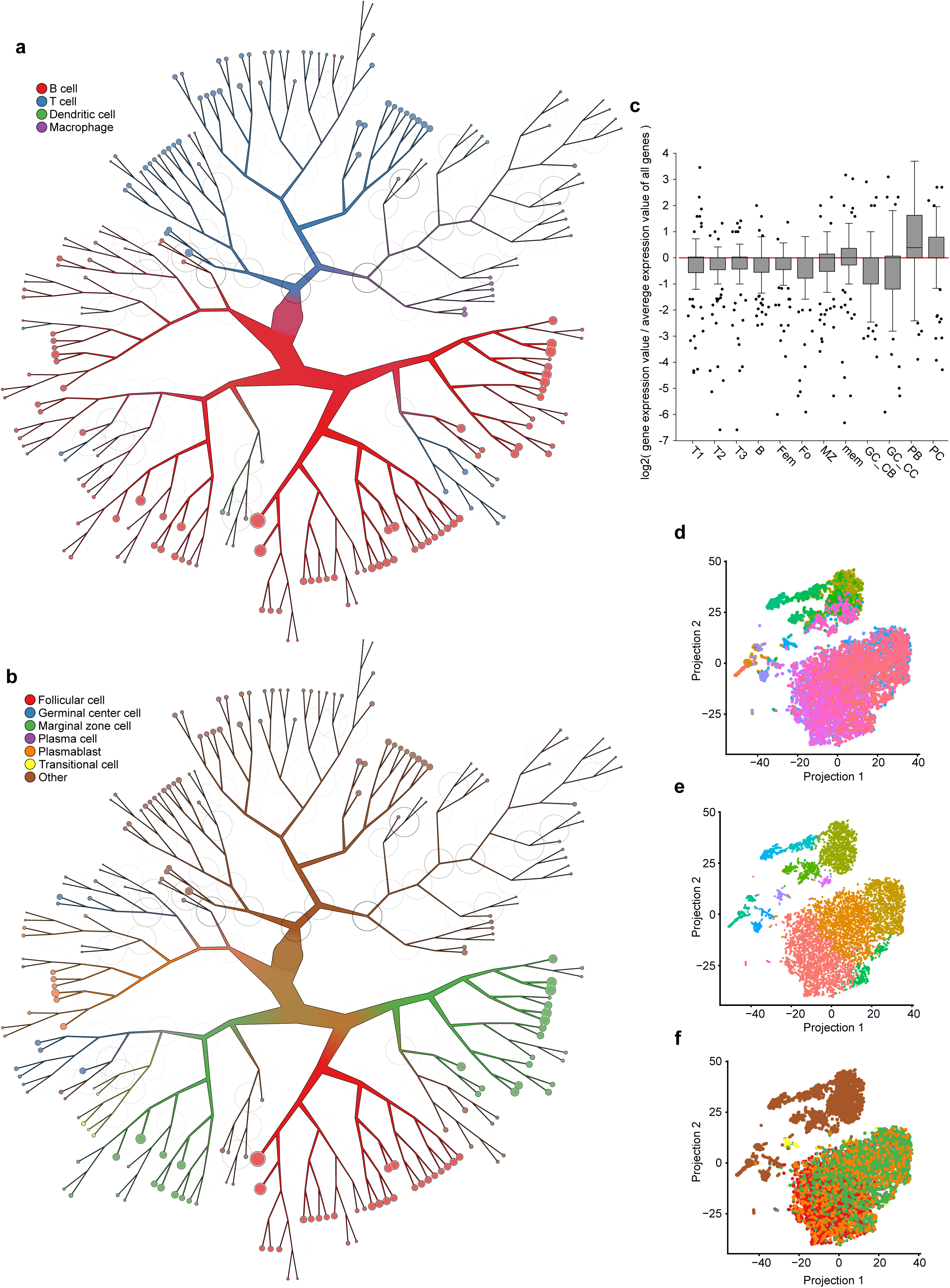
Plasmablast subpopulation identification in mouse spleen. (a) `TooManyCells` clustering tree of the mouse spleen data set labeled with previously identified cell types. (b) Tree from (a) with newly identified subtypes of B cells. (c) MyGeneSet gene expressions for the top 100 differentially expressed genes of the plasmablast node from (b) compared to all other cells. (d) Cells from (a) projected using Seurat’s processing and t-SNE, colored by `TooManyCells` clustering tree leaves, where each leaf is assigned a different color. Similar colors represent nearby locations within the tree, for example pink and purple are closer in the tree than pink and green. (e) Coordinates from t-SNE projection in (d) colored by subset populations from (b). Plasmablasts are orange and spread throughout the B cell grouping. (f) Coordinates from the t-SNE projection in (d) colored by Seurat-generated cluster labels.

To compare the identified plasmablasts to their location within a t-SNE projection, we used Seurat to generate t-SNE plots and clusters. Overlaying cells in the t-SNE projection with their leaves from the `TooManyCells` tree (Figure 5b), we observed an agreement on many cell locations – cells nearby in the projection were nearby in the tree (Figure 5d). However, there were some discrepancies where cells farther apart in the tree were nearby in the projection. To see whether the B cell subtypes were affected by this projection, we overlayed the B cell subtypes as defined by ImmGen^17^ from `TooManyCells` assignments (Figure 5b) onto the t-SNE coordinates generated by Seurat (Figure 5d). We found relative agreement with the locations of transitional, marginal zone, germinal center, and plasma cells (Figure 5e). However, the plasmablast population was distributed throughout the B cell population. Furthermore, Seurat’s clustering was also unable to identify a plasmablast cluster (Figure 5f). Together, these results show the benefit of hierarchical visualization and clustering in detection of rare populations from scRNA-seq.

## Discussion

Popular single cell clustering and visualizations have been firmly set in variations of flat clustering and t-SNE projections. While these methods are inherently useful for single cell analyses, they may be unsuitable for certain applications as demonstrated in this study. Here, we developed `TooManyCells` which provides alternative algorithms from clustering to visualization. `TooManyCells` uses a recursive technique to repeatedly identify subpopulations whose relationships are maintained in a tree. Compared to t-SNE, `TooManyCells` provides a fundamentally different visualization paradigm and enables a flexible platform for cell state stratification and exploration. `TooManyCells` provides an array of visualization features, a subset of them were presented here. In addition to clustering and visualization, `TooManyCells` provides several other capabilities including heterogeneity assessment, clumpiness that is a new way of quantifying relationships in single cell measurements given a set of attributes, and diversity and rarefaction statistics which were borrowed from ecology. A number of measures were used to benchmark `TooManyCells` performance. Comparison of `TooManyCells` to several commonly used algorithms demonstrated its effectiveness to both generate multiple layers of expected partitions and identify rare populations on several different subsets of 11 mouse organs.

Flexibility and versatility were considered in the `TooManyCells` design. `TooManyCells` is a generic framework consist of several algorithms that may be interchanged with other existing algorithms. The `TooManyCells` clustering module, `ClusterTree`, can be easily used for clustering of data types beyond scRNA-seq. `BirchBeer`, its visualization component, can be used to visualize any hierarchical structure. Together, our studies suggest further improvement of clustering and visualization techniques are needed to fully explore outputs of various single cell measurement technologies.

## Methods

### Clustering

`TooManyCells` clusters single cells using a matrix-free hierarchical spectral clustering process^3^. Spectral clustering using normalized cuts is a technique to partition data into groups, or clusters, where the items within a cluster are more similar to each other than they are to items within other clusters^18^. Unfortunately, this analysis is based on the pairwise similarity between items, leading to a computational complexity of *O*(*m*^2^) with *m* items^3^. Let **A** be a similarity matrix where **A**(*i, j*) represent the similarity between items *i* and *j* and **D**(*i, i*) = diag(**A1**) be the diagonal matrix where **1** is a column vector of 1’s. Briefly, let

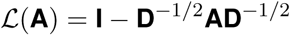

be the normalized Laplacian of **A**. A partition into two clusters denoted by 0 and 1 labels can be defined as

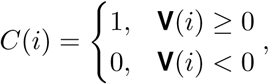

where **V** is the eigenvector corresponding to the second smallest eigenvalue of ℒ (**A \**)^18^. Alternatively, the eigenvector corresponding to the second largest eigenvalue of the shifted Laplacian,

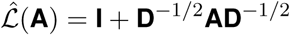

can be used instead of the second smallest eigenvalue of Laplacian matrix. Optionally, `TooManyCells` can consider additional eigenvectors using kmeans clustering^19^. This process bipartitions the data into two clusters. However, this is a slow process and computationally impractical as the number of measured single cells increases. To improve the speed of spectral clustering while retaining the original accuracy, `TooManyCells` implements a generalized version of algorithm that was originally proposed in^3^ for text mining for for use with sparse scRNA-seq matrices or any other observation / feature matrix. This implementation circumvents explicitly calculate of **A** and SVD.

To this end, let **B**_1_ be an *m* × *n* matrix with *m* rows of cells and *n* columns of read counts. `TooManyCells` takes as input a transpose of this matrix to conform to the current single cell matrix file format standards where the cells are c olumns. `TooManyCells` offers as by default the option to remove columns (genes) with no reads and rows (cells) with < 250 read counts. Then, for all 1 ≤ *i* ≤ *m,* 1 ≤ *j* ≤ *n*,

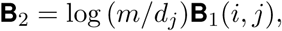

where 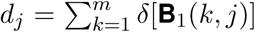 and *δ*(*x*) is 1 if *x* ≥ 1 and 0 if *x* = 0, for all *x* ∈ ℤ^+^. This normalization transformers **B**_1_ into a term frequency-inverse document frequency (tf-idf) matrix **B**_2_^13,14^, where the importance of common genes are de-emphasized for clustering. Intuitively, a ubiquitously expressed gene is unlikely to be as important for cell clustering compared to a gene only expressed in a given subpopulation. Other data normalizations can be performed prior to this transformation or replace the tf-idf process entirely. For instance, one may normalize each cell based on its total read count followed by the normalization of each gene by that gene’s median positive read count. In order to relate cells in a matrix-free manner, cosine similarity was used^20^. It has been shown that the similarity matrix **A** can be derived in terms of as^3^

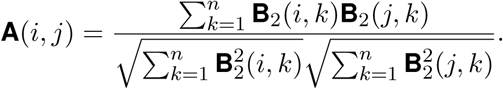

However, in order to lower the computational complexity, `TooManyCells` does not calculate this matrix. Instead, a new matrix **B** is defined as

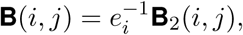

where 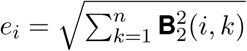 is the Euclidean norm of **B**_2_ row *i*.

To prepare the matrix as a form of a normalized Laplacian, let

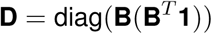

and

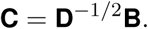

Then the eigenvector of ℒ (**A**) corresponding to the second smallest eigenvalue is the second left singular vector corresponding to the second largest singular value of **C**, which can be found using truncated singular vector decomposition (SVD)^3^. It has been shown that the computation complexity of this process is *O*(*Jm*), the number of non-zero entries of **C**, where *J* is the average number of expressed genes within a cell.

This bipartition can be recursively applied to each delineated cluster until a stopping criteria is reached. This recursive clustering results in a divisive hierarchical cluster structure. In accordance with^3^, `TooManyCells` uses Newman-Girvain modularity (*Q*)^4^. Modularity is a measure from community detection which has also been used in single cell clustering through optimization using the Louvain method^1,11,12^. Let *G* = (*V, E*) be a weighted graph of *m* nodes (cells) with *e* edges. Newman-Girvain modularity measures the strength of the partition of nodes. For a bipartition,

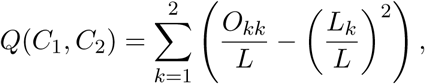

where 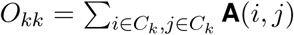 **A**(*i, j*) is the total degree of nodes in cluster *C*_*k*_, if 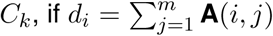 is the degree of node *i* then 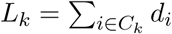 is the total degree of nodes in *C*_*k*_, and 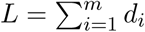 is the degree of all nodes in the network. *Q* measures the distance of edges within clusters to the random distribution of clusters, such that *Q* > 0 denotes non-random communities and *Q* ≤ 0 demonstrates communities randomly found^3,4^.

`TooManyCells` uses *Q* to assess a candidate partition of cells to determine whether to continue the recursion or stop as a leaf in the divisive hierarchical clustering. That is, at each bipartion, if *Q* > 0 then continue the recursion, otherwise stop. Thus, the end result of this top-down clustering is a tree structure of clusters, where each inner node is a clustering and the leaves are the most reliable fine grain clustering where any additional splitting would lead to random partitioning of cells. This process has *O*(*Jm* log *m*) computational complexity^3^. The code for the `TooManyCells` implementation of this algorithm is available at https://github.com/faryabiLab/too-many-cells.

### Visualization

The `TooManyCells` clustering algorithm results in a tree structure, where each inner node is a coarse cluster and each leaf is the most refined cluster per modularity measure. The `BirchBeer` rendering method was developed for displaying single cells cluster hierarchies. `BirchBeer` was developed specifically to address visualization and interpretation of single-cell resolution displays of scRNA-seq data. To this end, `BirchBeer` utilizes `graphviz` for node coordinate placement and the Haskell `diagrams` library as rendering engine.

`BirchBeer` provides a multitude of graphical features to assist in the detection and interpretation of cell clusters. The tree leaves can be displayed in various ways. Single-cell resolution exploration is facilitated by drawing color-coded individual cells at the tree leaves. Alternatively, a pie chart can be shown to visualize a summary of the cell composition of the clusters at the tree leaves. Both single-cell resolution and statistical summarization can be shown using a “pie ring”. Each tree branch can be scaled to the relative number of cells within each subtree, allowing for quick inspection of cell population sizes of various clustering levels and visualizing clusters of rare and common populations. Furthermore, colors can be applied to each branch such that the weighted average blend of the colors of each label in the subtree is used, allowing for immediate detection of subtrees with large differences or similarities. Cluster numbers can be displayed on each node, tracing the data back into a human readable interpretation of differences between the clusters at various hierarchy levels. Furthermore, the modularity of each candidate split can be displayed at each node as a black circle with varying darkness to demonstrate the dissimilarity of cell populations encompassing that assay. Large trees may result in busy figures, much like large t-SNE plots, so options to prune the tree are available. Cutting the tree at certain levels, node sizes, or modularity are some options, but additionally there is a statistically driven option called `--smart-cutoff` which cuts the tree depending on the median absolute deviation (MAD). For instance, a stopping criteria of four MADs from the median node size to keep the structure of the tree but prune smaller branches. `BirchBeer` accepts JSON trees as a standard input. The code for `BirchBeer` is available at https://github.com/faryabiLab/birch-beer.

### Differential expression

Given multiple cluster identification numbers, `TooManyCells` can perform differential expression analysis to identify the difference between the gene expression of cells in these clusters. `TooManyCells` uses `edgeR` for differential expression analysis^21^. To visually facilitate this analysis, `BirchBeer` can label clusters with their identification numbers.

### Diversity analysis

While Shannon entropy^7^ is frequently used as a measure of “diversity”, the effective number of species is a more meaningful measure of diversity in biological settings. For example, a population with 16 equally abundant species should be twice as diverse as a population with 8 equally abundant species^9^. Assuming each cell is an “organism” belonging to a “species” group defined by the clustering algorithm, then a diversity index can be applied to find the effective number of cell states in a population.

The diversity satisfying such a property can be defined as^8,9^

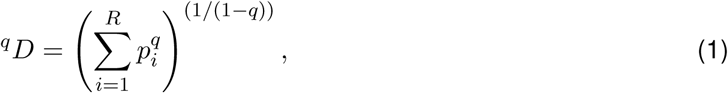

where *p*_*i*_ is the frequency of species *i, R* is the total number of species in the population, and *q* is the “order” of diversity. *q* > 1 gives additional weight towards common species, while more weight is given to rare species when *q* < 1. *q* = 1 gives equal weight to all the species regardless of their commonality and is defined as

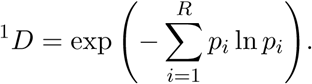

Several diversity measures can be derived from equation 1 For instance, ^0^*D* defines richness, or the number of species, in the population. ^1^*D* relates to c exp Shannon entropy and ^2^*D* is the inverse of the Simpson index^22^. Various diversity measures have been used previously in domains such as lymphocyte receptor repertoires and cell clones^23–25^. Here, we use the diversity in `TooManyCells` to quantitate the effective number of cell states within a population.

`TooManyCells` borrows the concept of rarefaction curve from ecology^26,27^ to estimate the number of detectable species in a given number of profiled single cells. Briefly, the estimated number of species in a population can be calculated from a given number of samples taken from a population through random subsampling. The estimated number of species in a subsample of size *n* representing *X*_*n*_ species can be calculated as

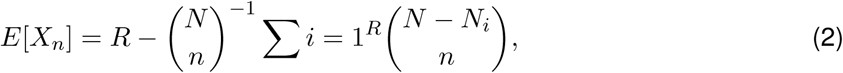

where *N* is the total number of cells, *R* is the total number of cell states in all samples, and *N*_*i*_ is the number of cells belonging to state *i*. For the interval [0, *R*], the equation 2 generates a rarefaction curve that shows the estimated number of species for a given number of profiled cells. The steepness of the rarefaction curve may represent the heterogeneity of a population. For a given number of subsamples, the estimated number of species across multiple populations can be compared based on their respective rarefaction curves. This property is useful for comparing populations with different sample sizes. A plateau in the curves indicates no substantial increase in the number of new cell states, implying a sufficient sampling to observe all the cell states in a sample. `TooManyCells` implements this procedure to rarefy populations.

### Clumpiness

The hierarchical structure generated from any hierarchical clustering, both divisive and agglomerative, holds items in the leaf nodes. Each item can be assigned a label, such as a tissue of origin, cell type, or expression level of high or low, where the items here are cells. In order to quantify the level of aggregation within the tree, a measure of “clumpiness” is needed^15^. For instance, the degree of how “clumped”, or co-localized, are CD4 T cells and CD8 T cells within the tree. Here, one would expect those T cells to be grouped together more closely than CD4 T cells with B cells. A clumpiness measure enabled the quantification of this similarity.

The clumpiness measure used here was specifically designed for hierarchical structures and described in more detail in^15^. Briefly, consider a rooted *k* -*ary* tree. The clumpiness of the set of leaves *M* when partitioned according to *L* = {*L*_1_, *L*_2_, *…, L*_*n*_} is defined as

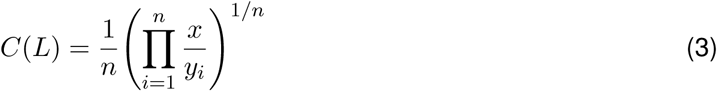

This measure takes the geometric mean of *x* weighted by the frequency of each label, *y*_*i*_. *x* represents the weighted number (weighted by distance to the descendant leaves) of “viable” non-root inner nodes weighted by *y*_*i*_. Viable nodes are comprised of inner nodes that have at least one vertex of each label in their descendant leaves. The clumpiness of a label *L*_*i*_ with itself is simply considering an *L*′ containing two sets — leaves in *L*_*i*_ and all other leaves. Then the clumpiness of *L*_*i*_ with itself is 1 - *C*(*L*′)^15^.

### Data availability

Microfluidics single cell RNA-seq count data from 11 organs in 3 female and 4 male, C57BL/6 NIA, three month old mice were obtained from https://figshare.com/articles/_/5715025, removing P8 libraries due to outlier cell counts^6^.

**Figure S1:**
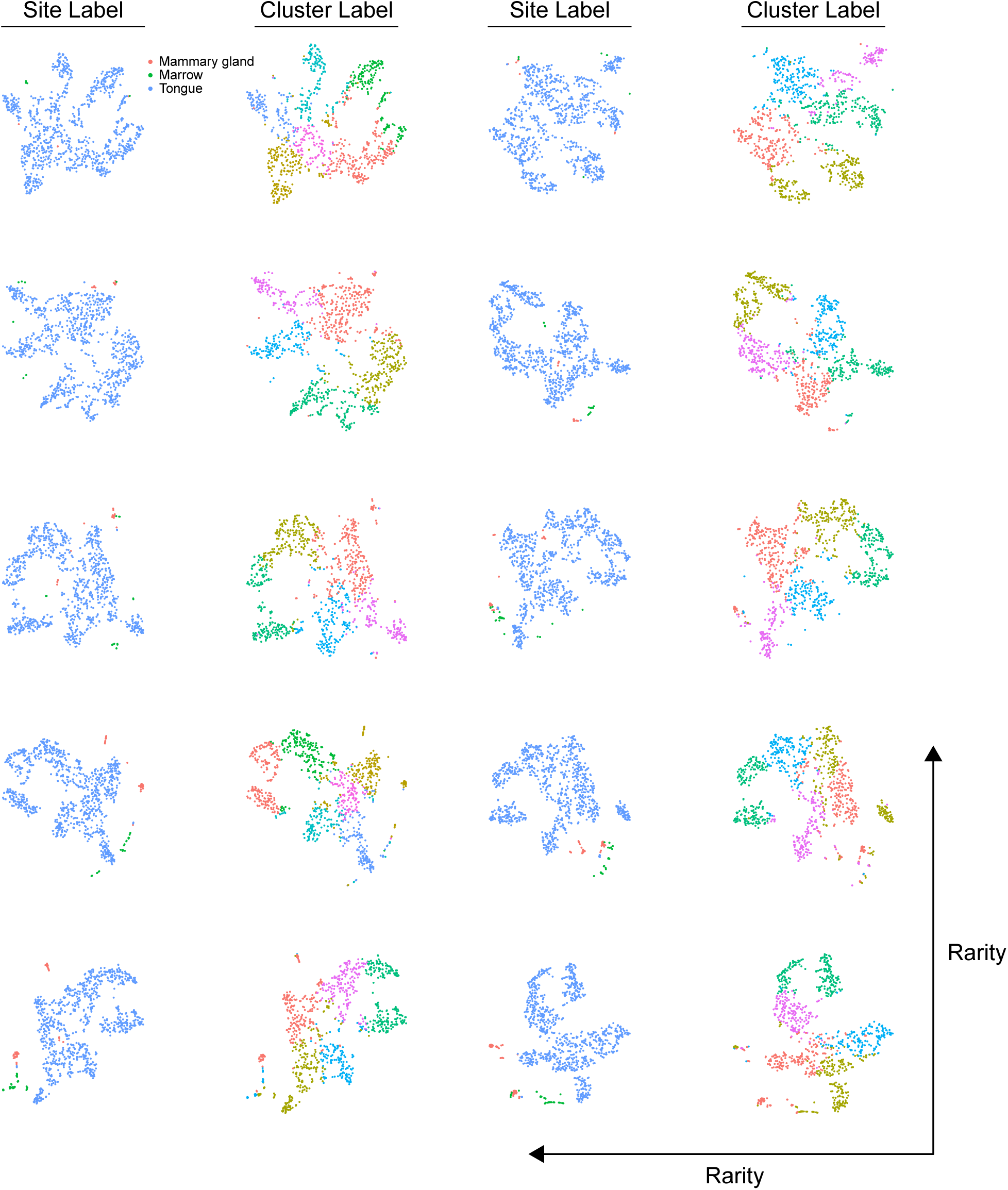
t-SNE projections of the rare population benchmark using Cellranger. Increasing rare population from 10 (top left) to 100 (bottom right).

**Figure S2:**
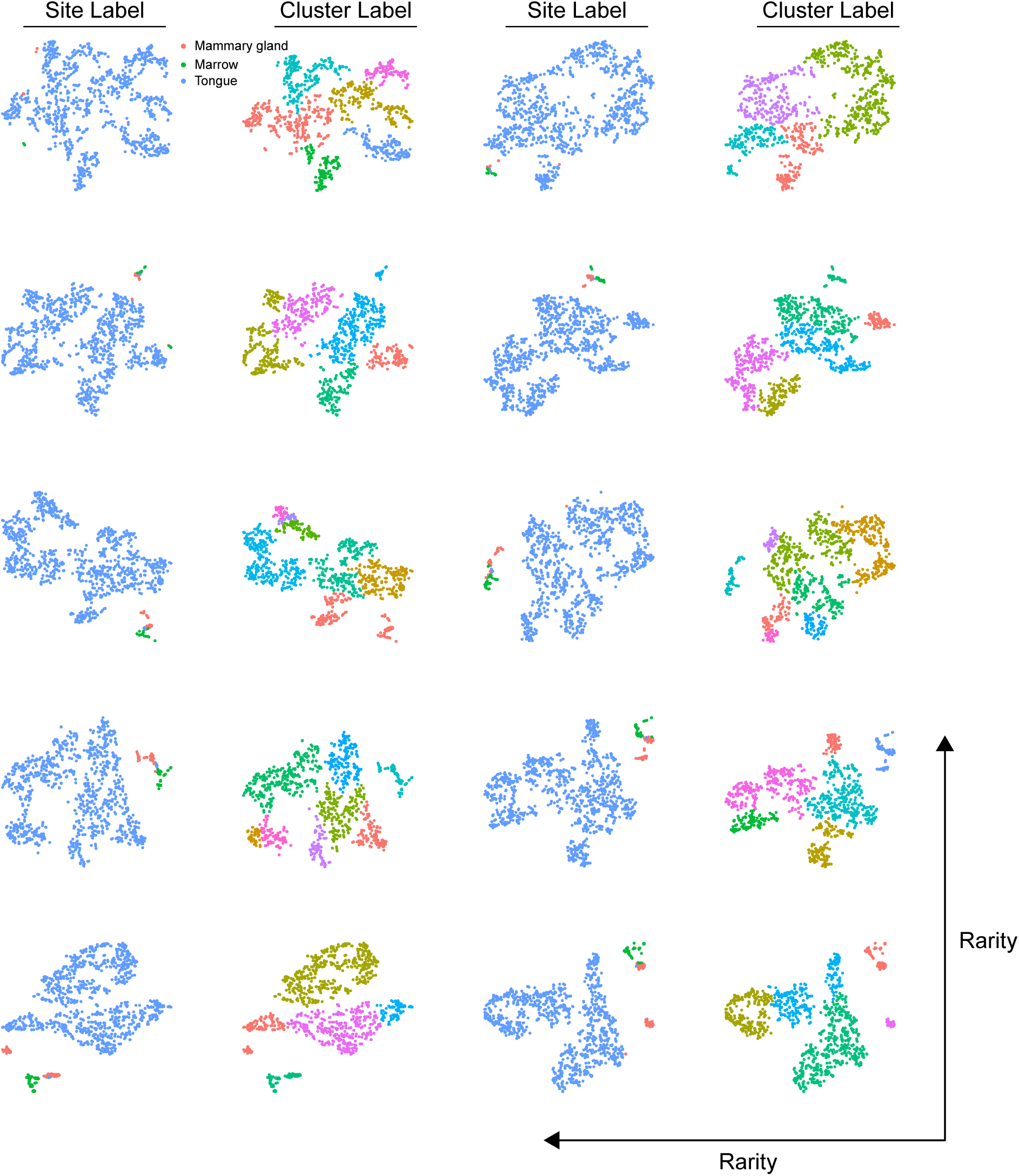
t-SNE projections of the rare population benchmark using Monocle. Increasing rare population from 10 (top left) to 100 (bottom right).

**Figure S3:**
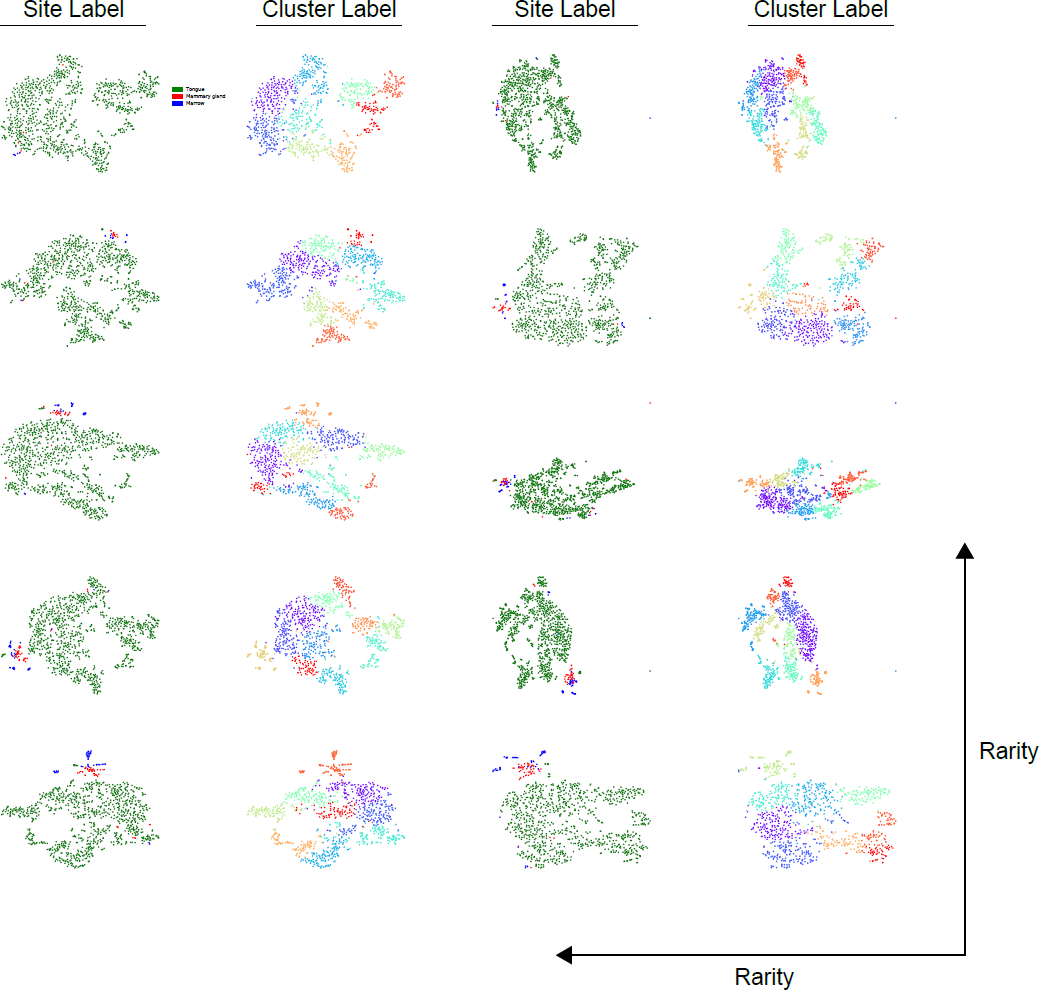
t-SNE projections of the rare population benchmark using Phenograph. Increasing rare population from 10 (top left) to 100 (bottom right).

**Figure S4:**
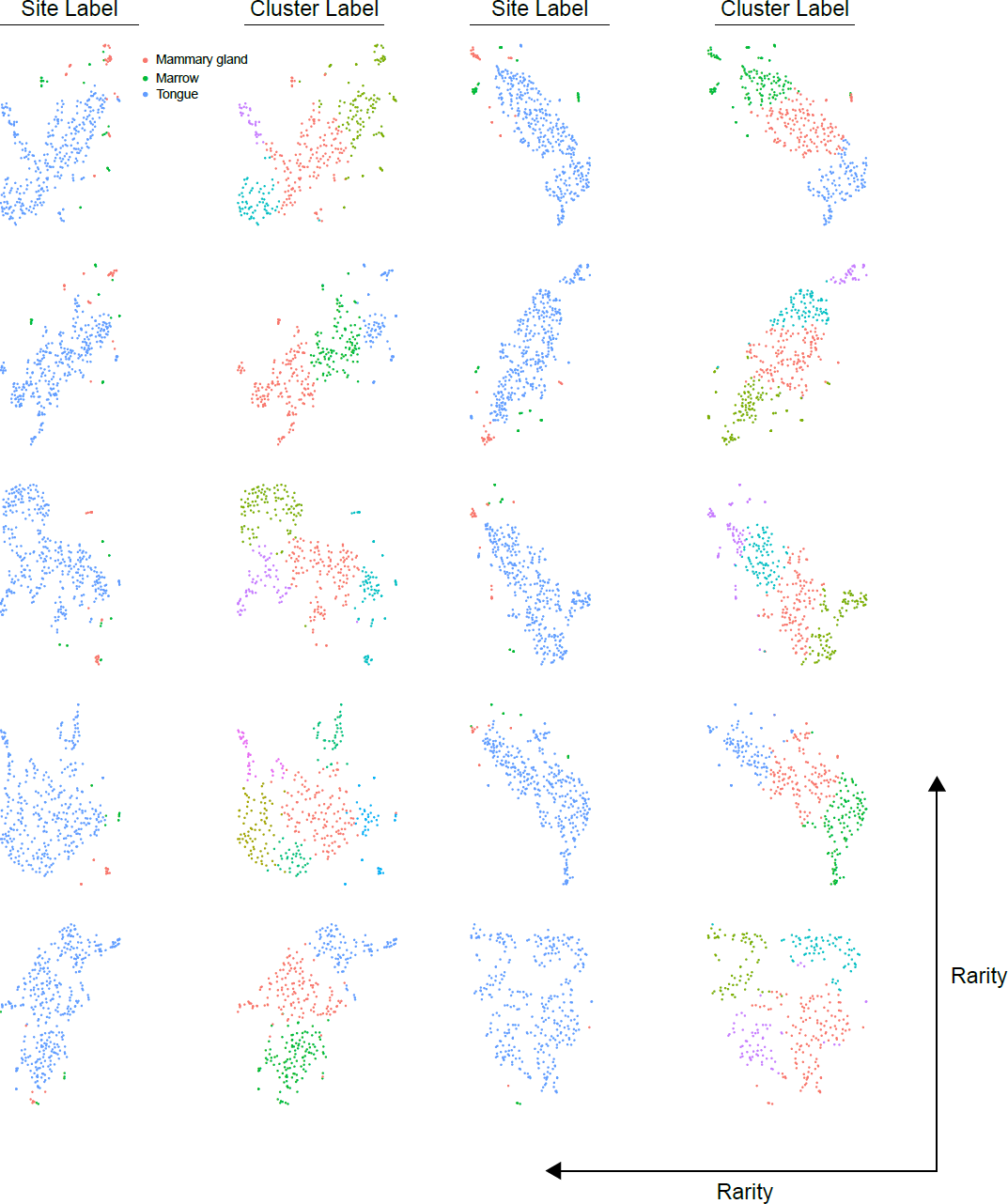
t-SNE projections of the rare population benchmark using Seurat. Increasing rare population from 10 (top left) to 100 (bottom right).

**Figure S5:**
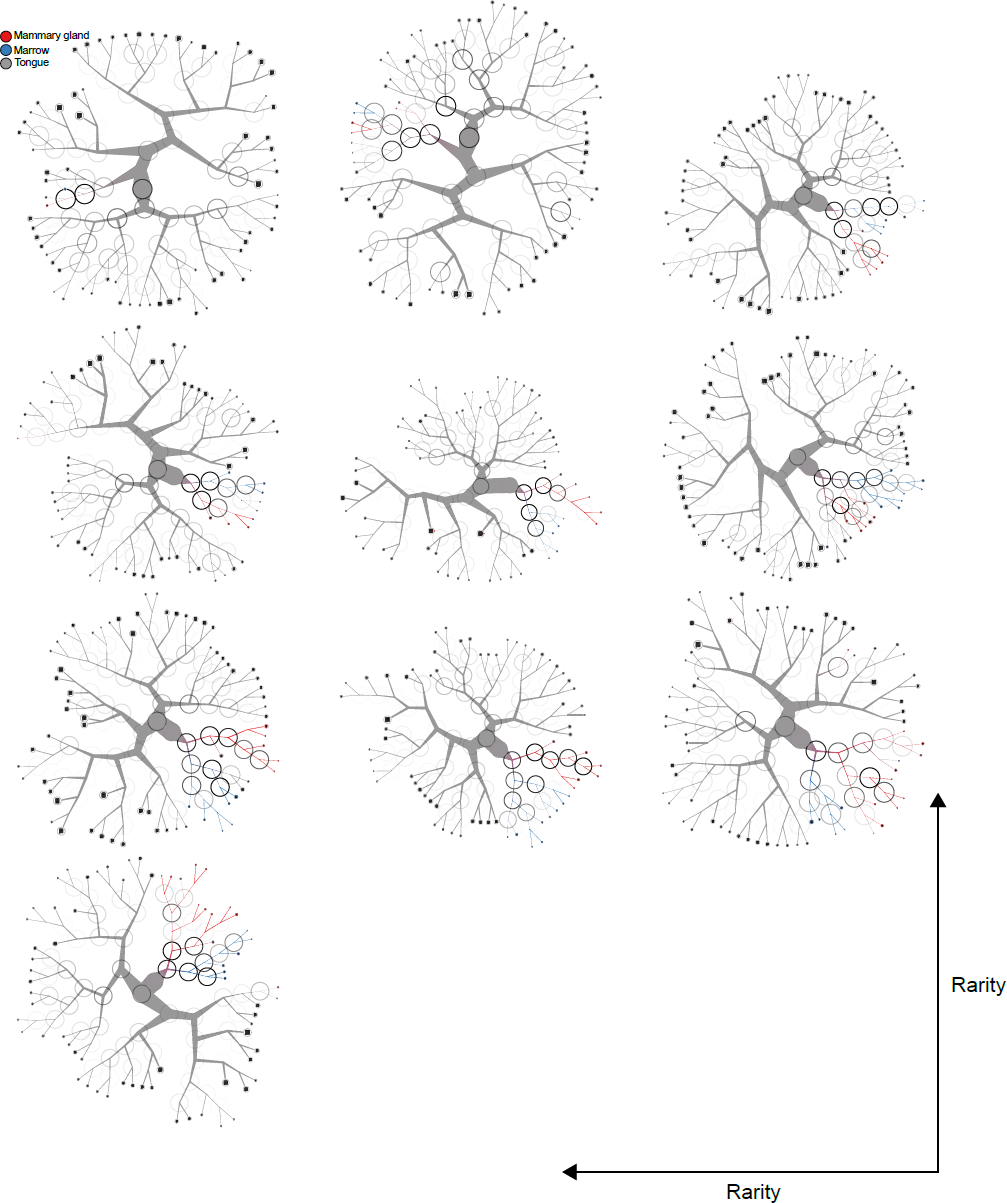
t-SNE projections of the rare population benchmark using `too-many-cells`. Increasing rare population from 10 (top left) to 100 (bottom right).

**Figure S6:**
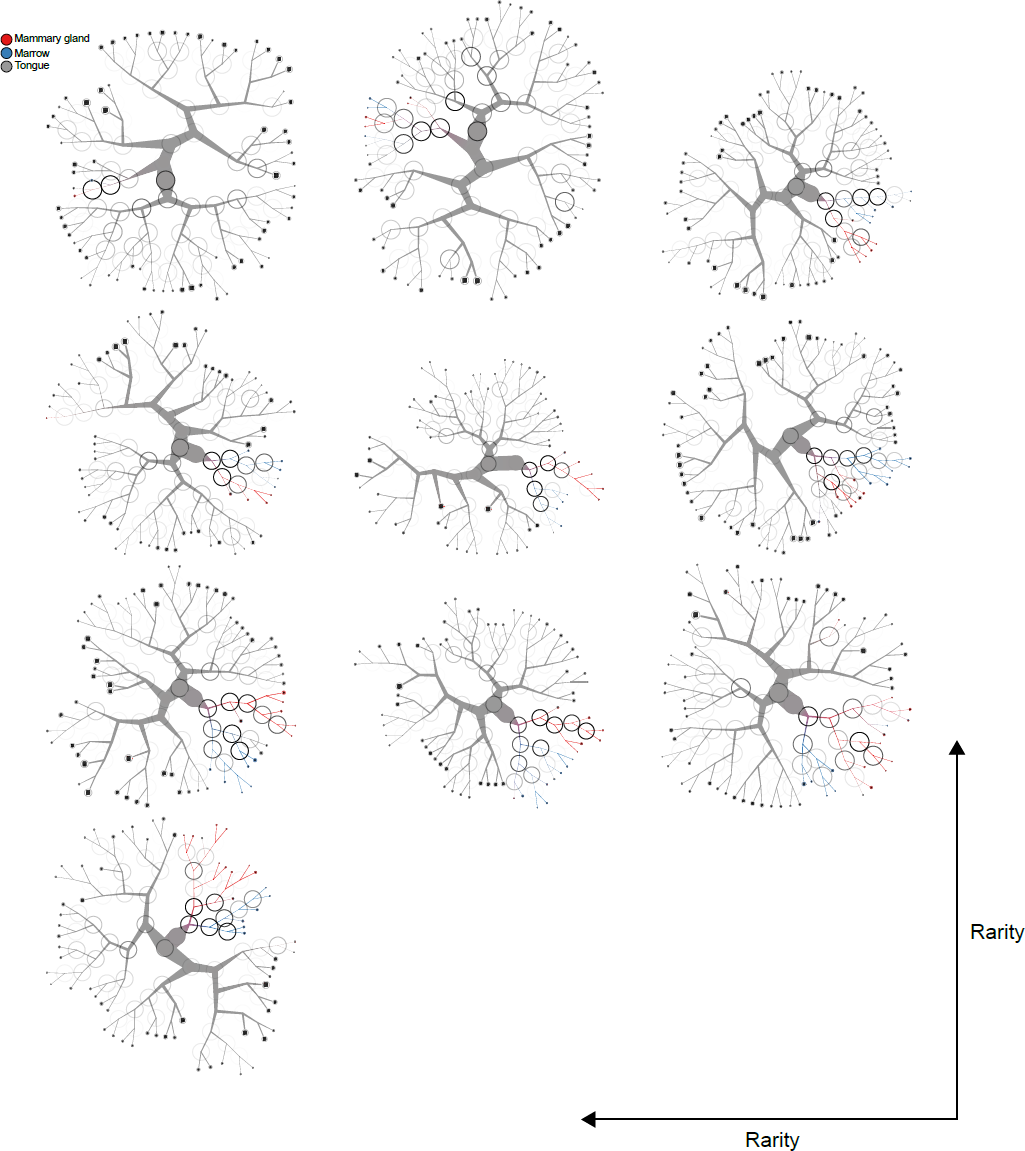
t-SNE projections of the rare population benchmark using `too-many-cells` with both normalization methods. Increasing rare population from 10 (top left) to 100 (bottom right).

**Figure S7:**
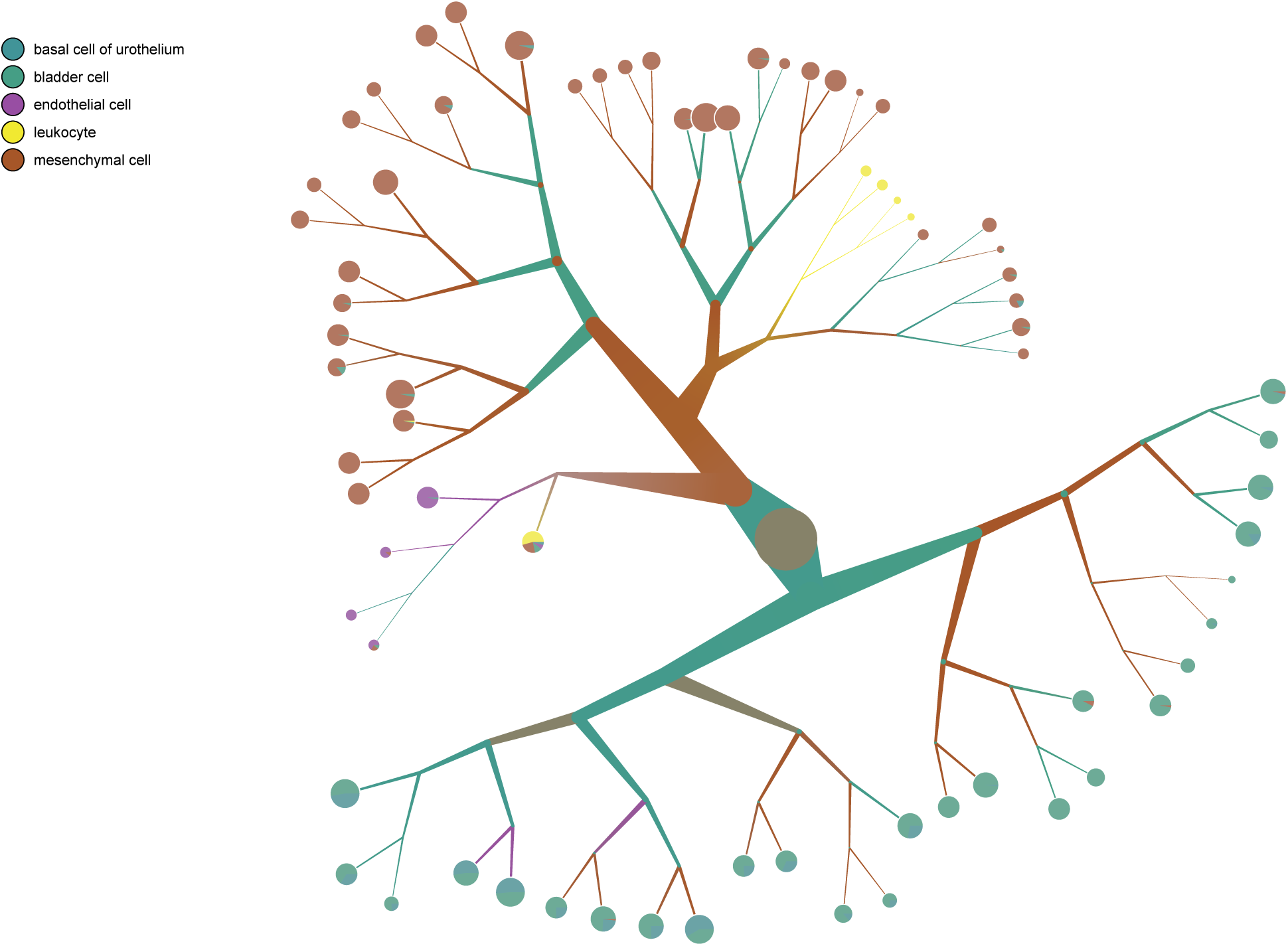
`TooManyCells` tree with cell type overlay in bladder.

**Figure S8:**
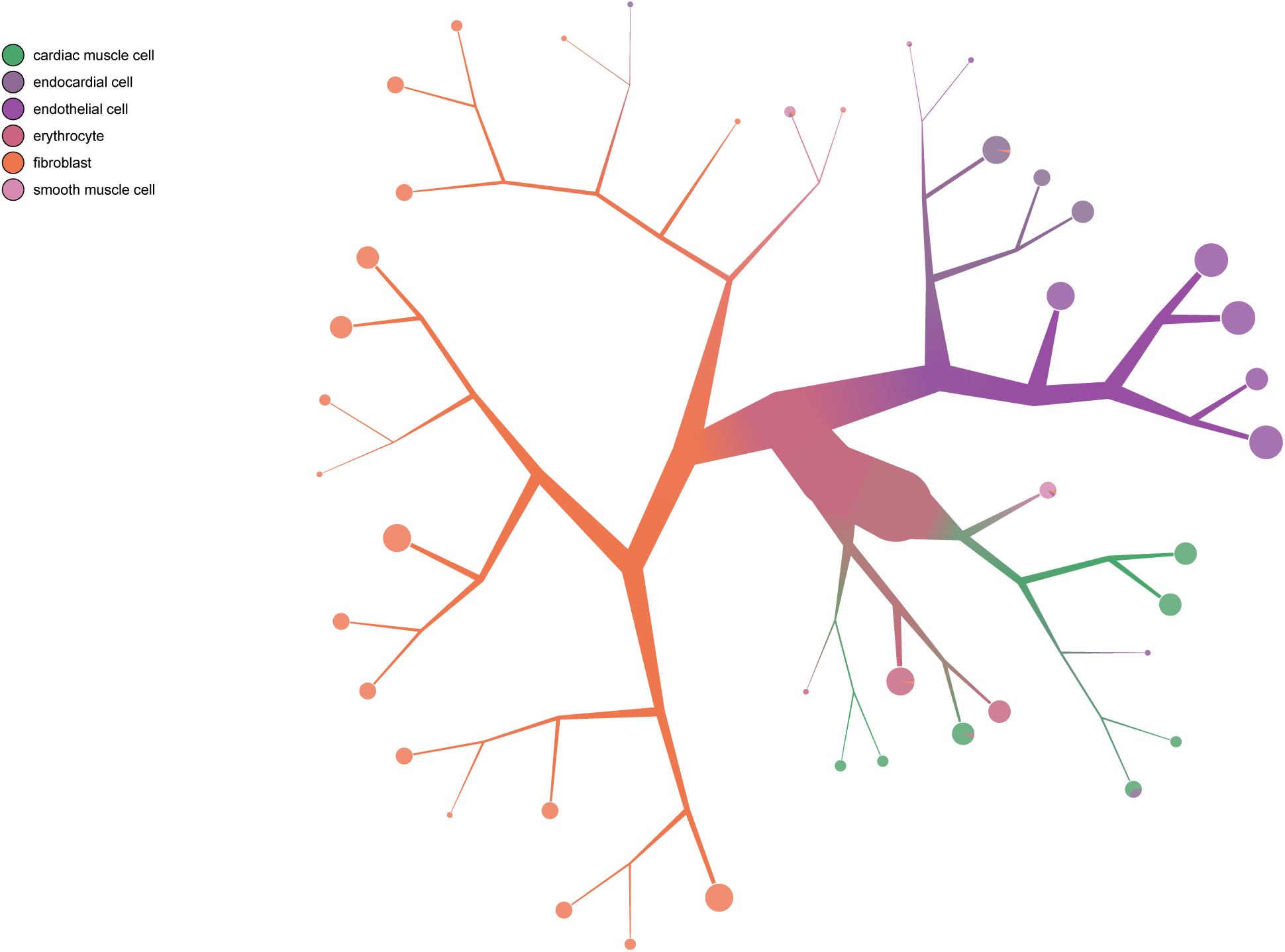
`TooManyCells` tree with cell type overlay in heart.

**Figure S9:**
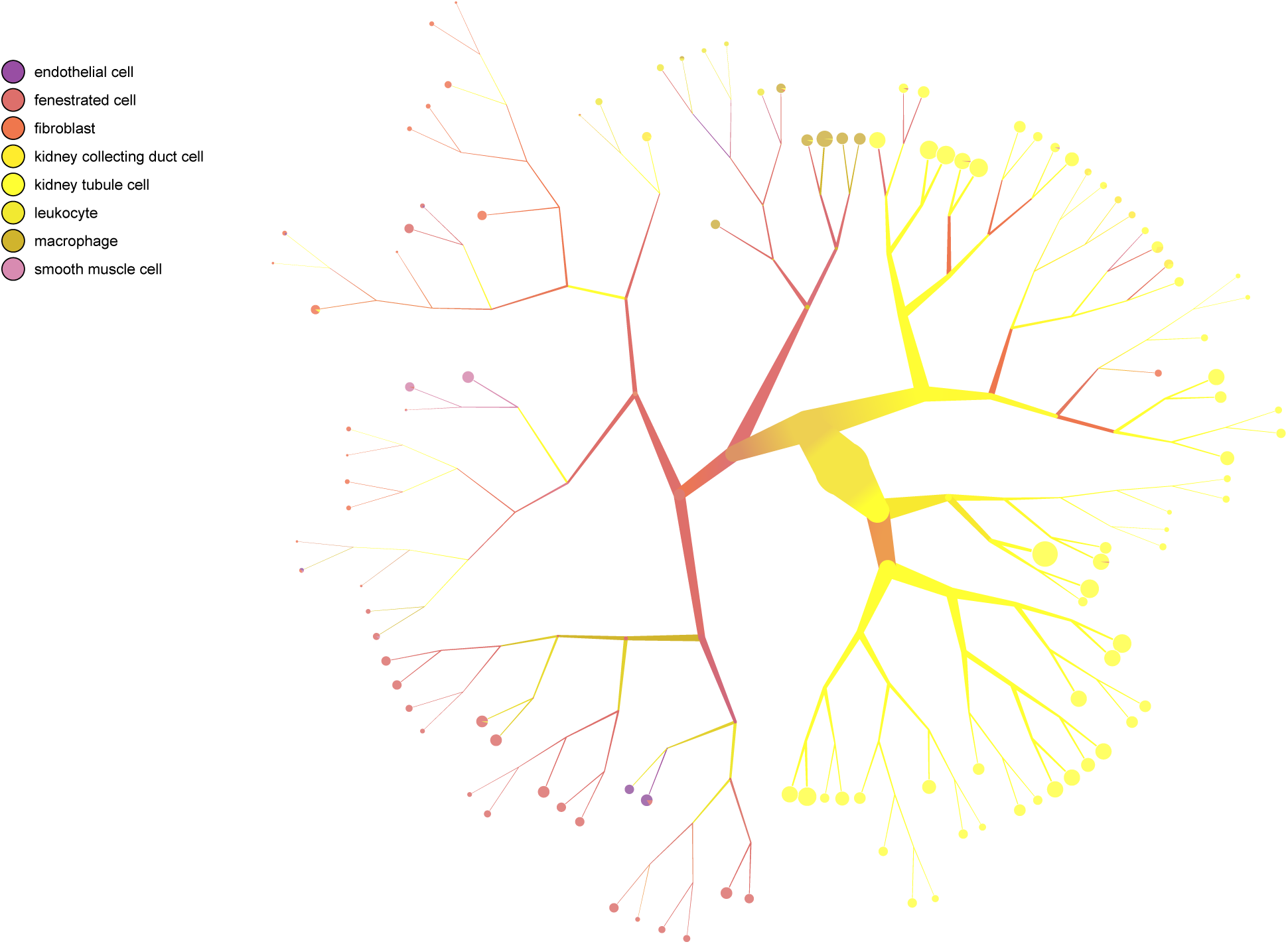
`TooManyCells` tree with cell type overlay in kidney.

**Figure S10:**
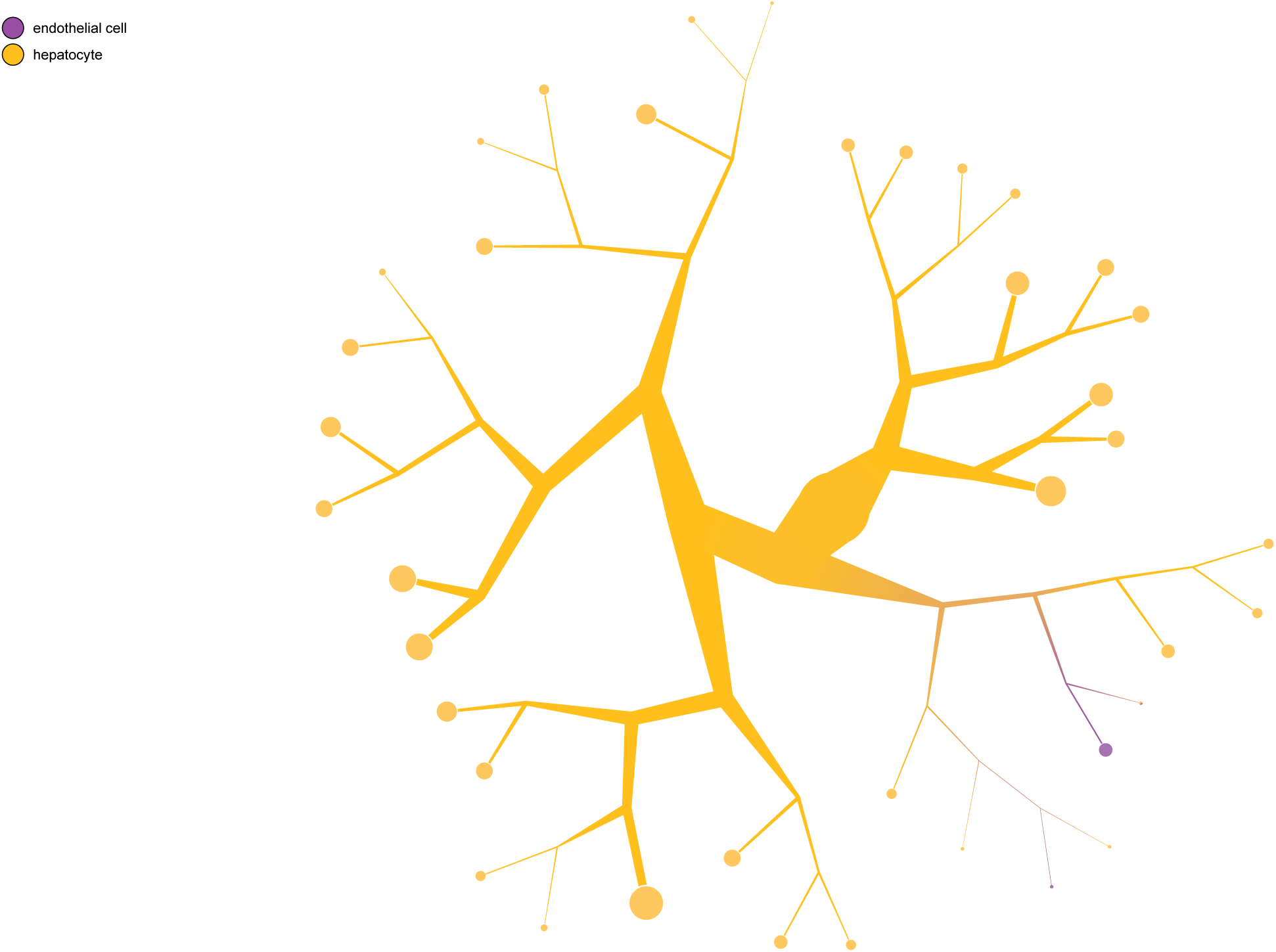
`TooManyCells` tree with cell type overlay in liver.

**Figure S11:**
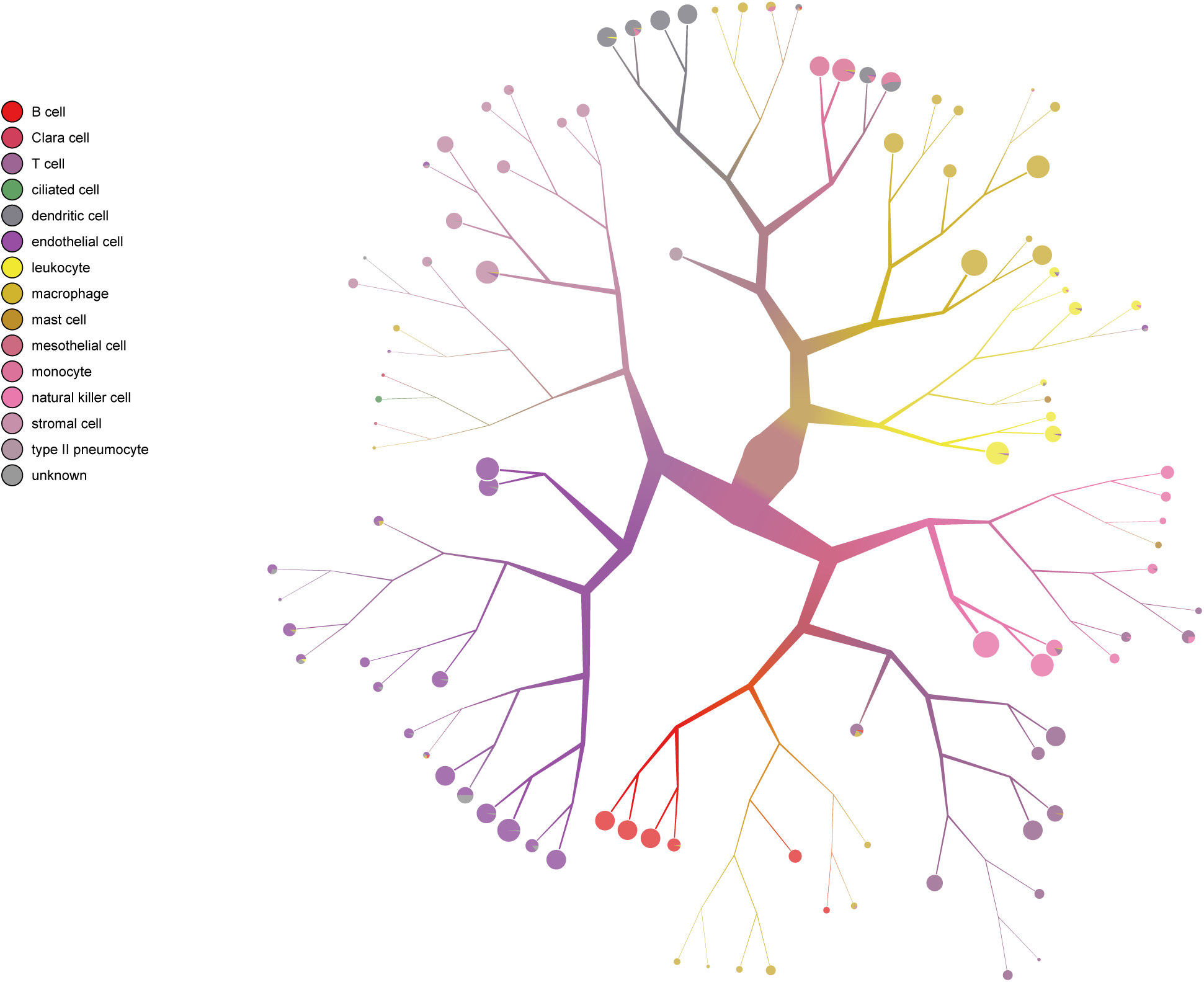
`TooManyCells` tree with cell type overlay in lung.

**Figure S12:**
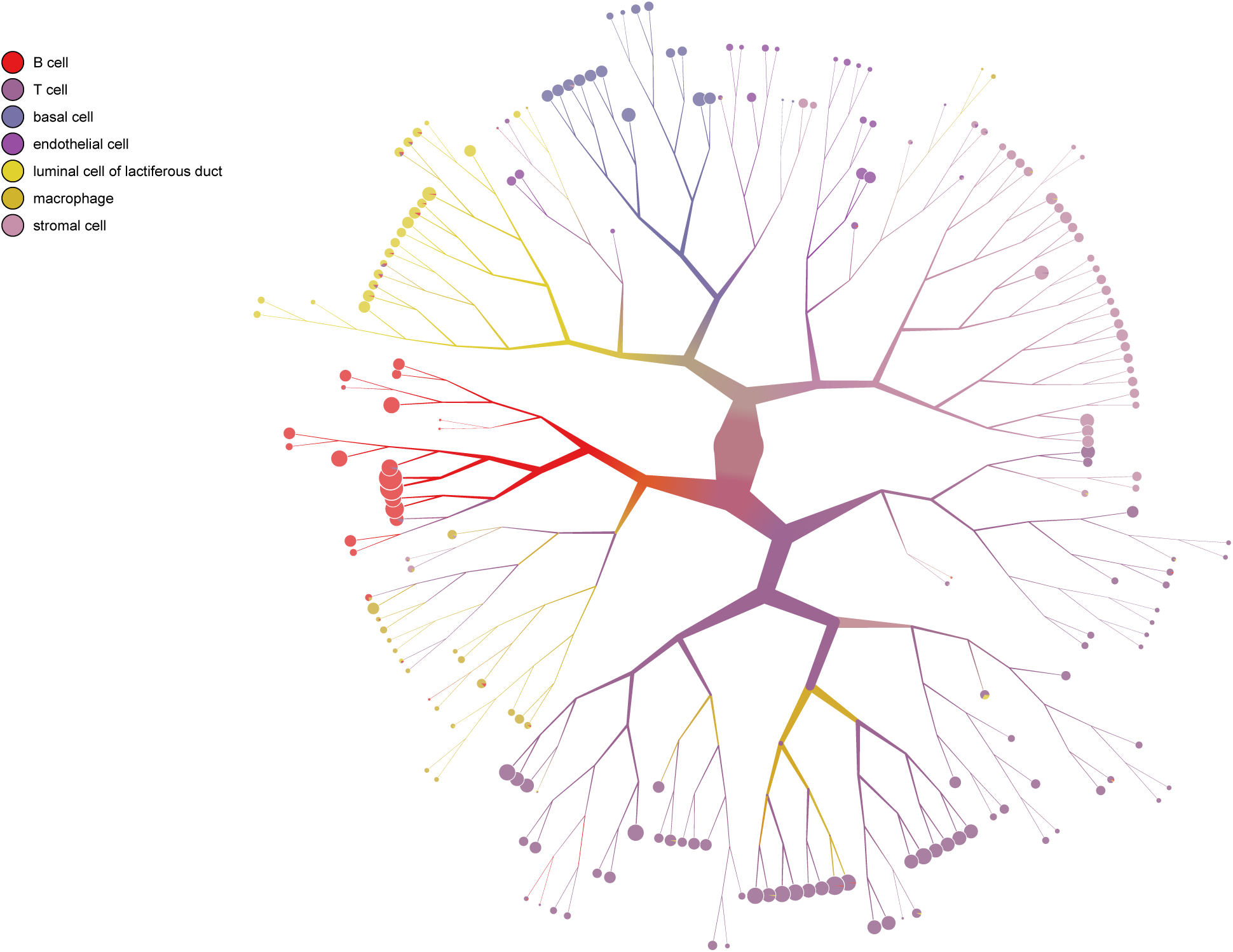
`TooManyCells` tree with cell type overlay in mammary gland.

**Figure S13:**
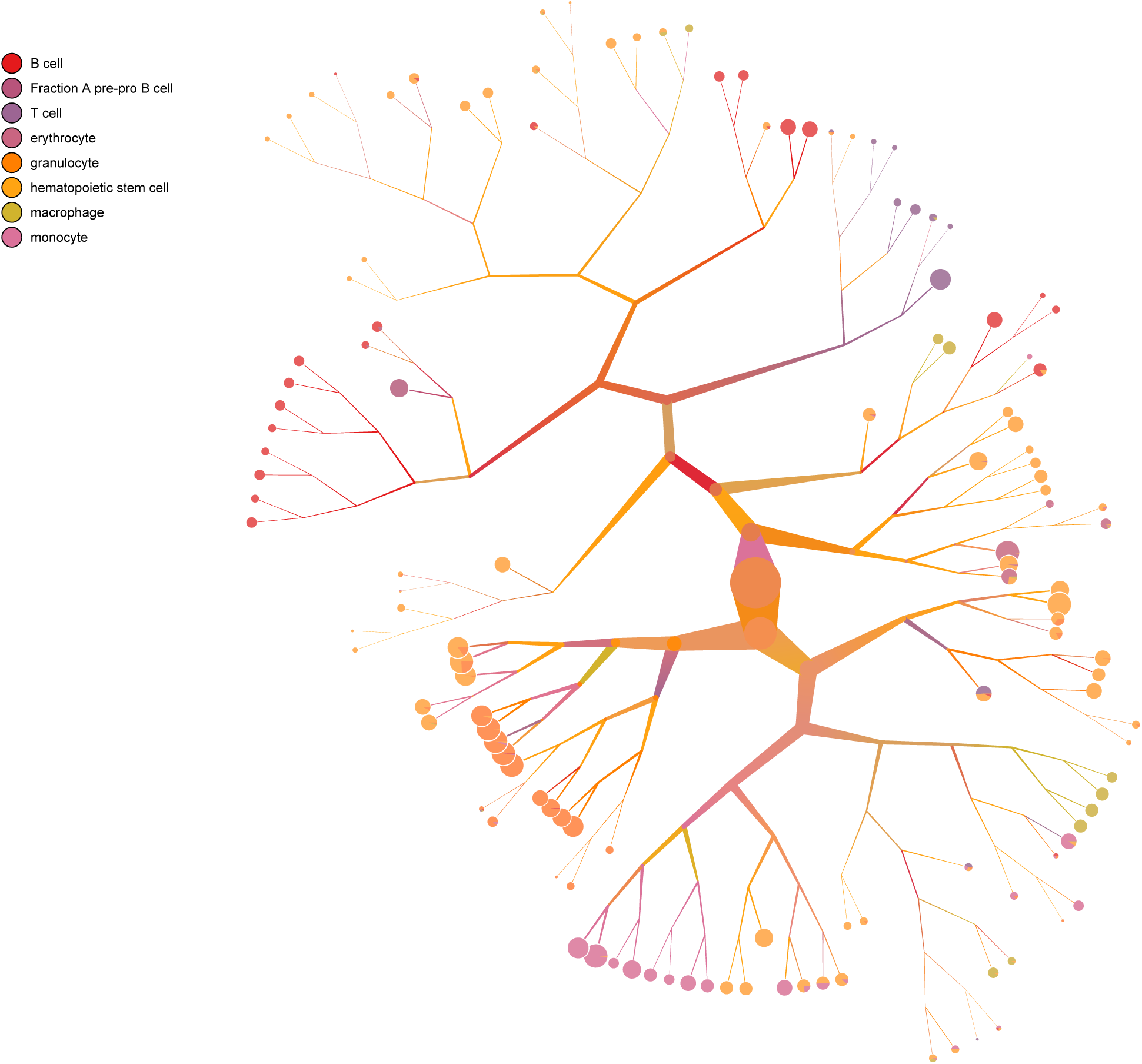
`TooManyCells` tree with cell type overlay in marrow.

**Figure S14:**
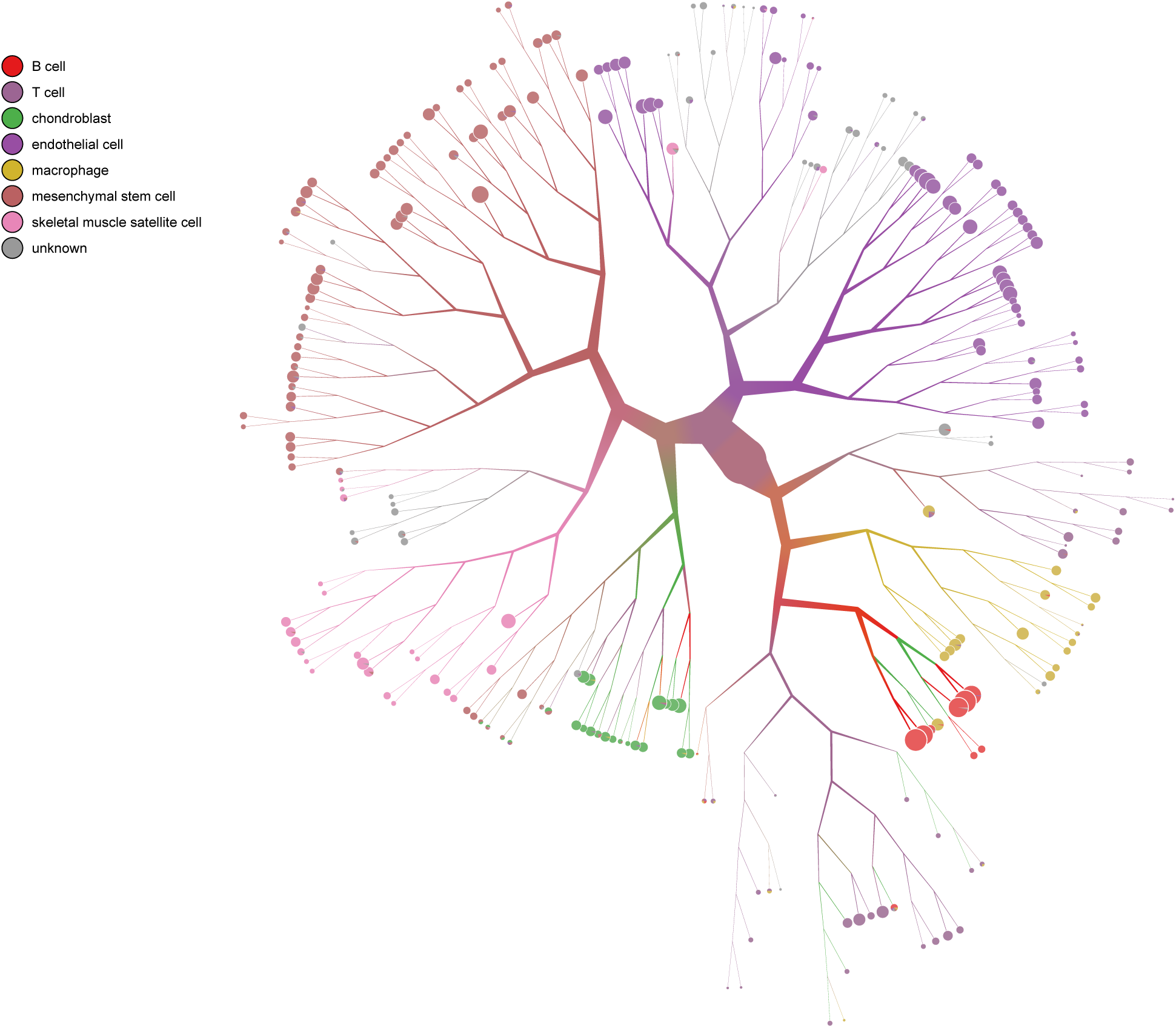
`TooManyCells` tree with cell type overlay in limb muscle.

**Figure S15:**
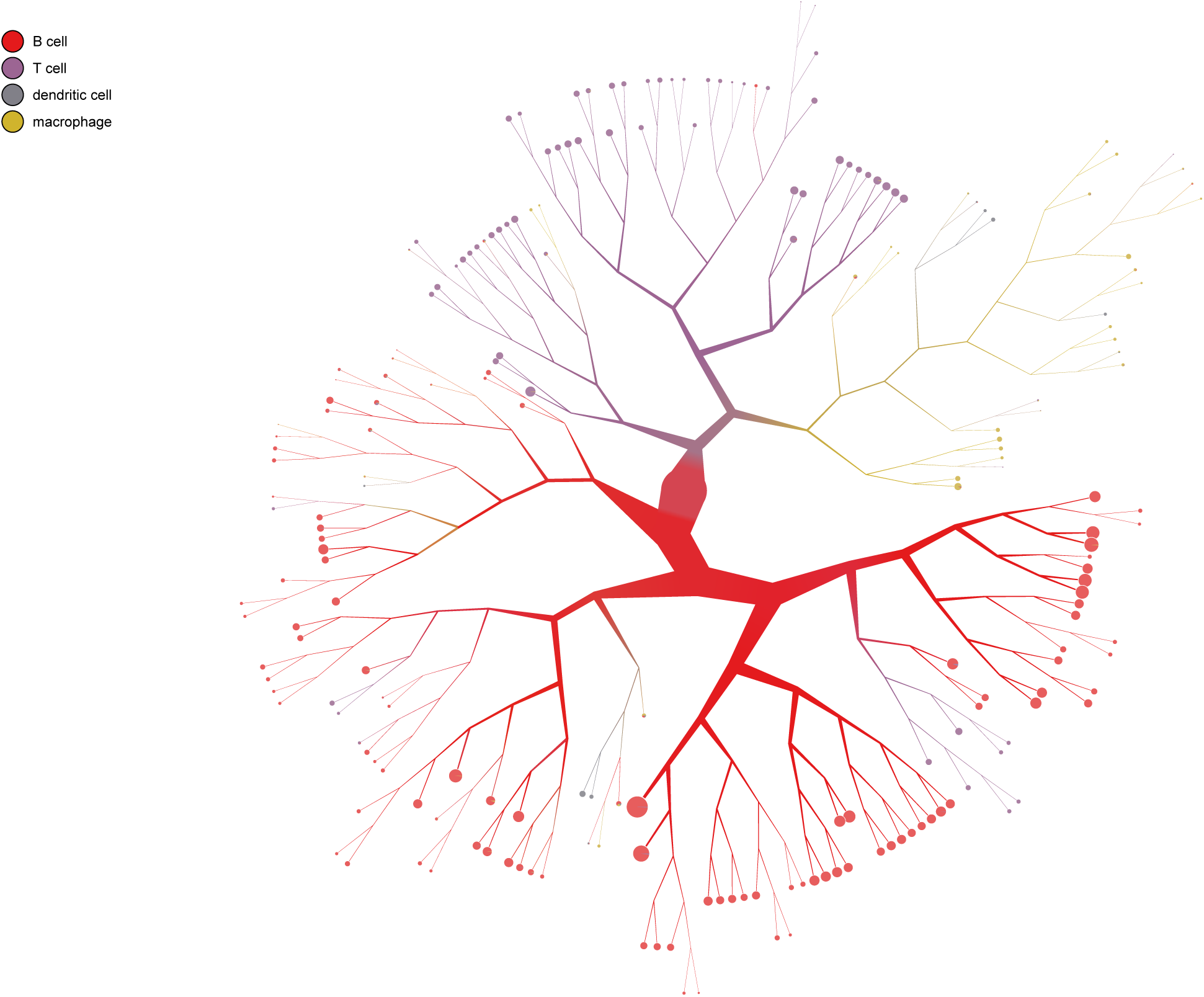
`TooManyCells` tree with cell type overlay in spleen.

**Figure S16:**
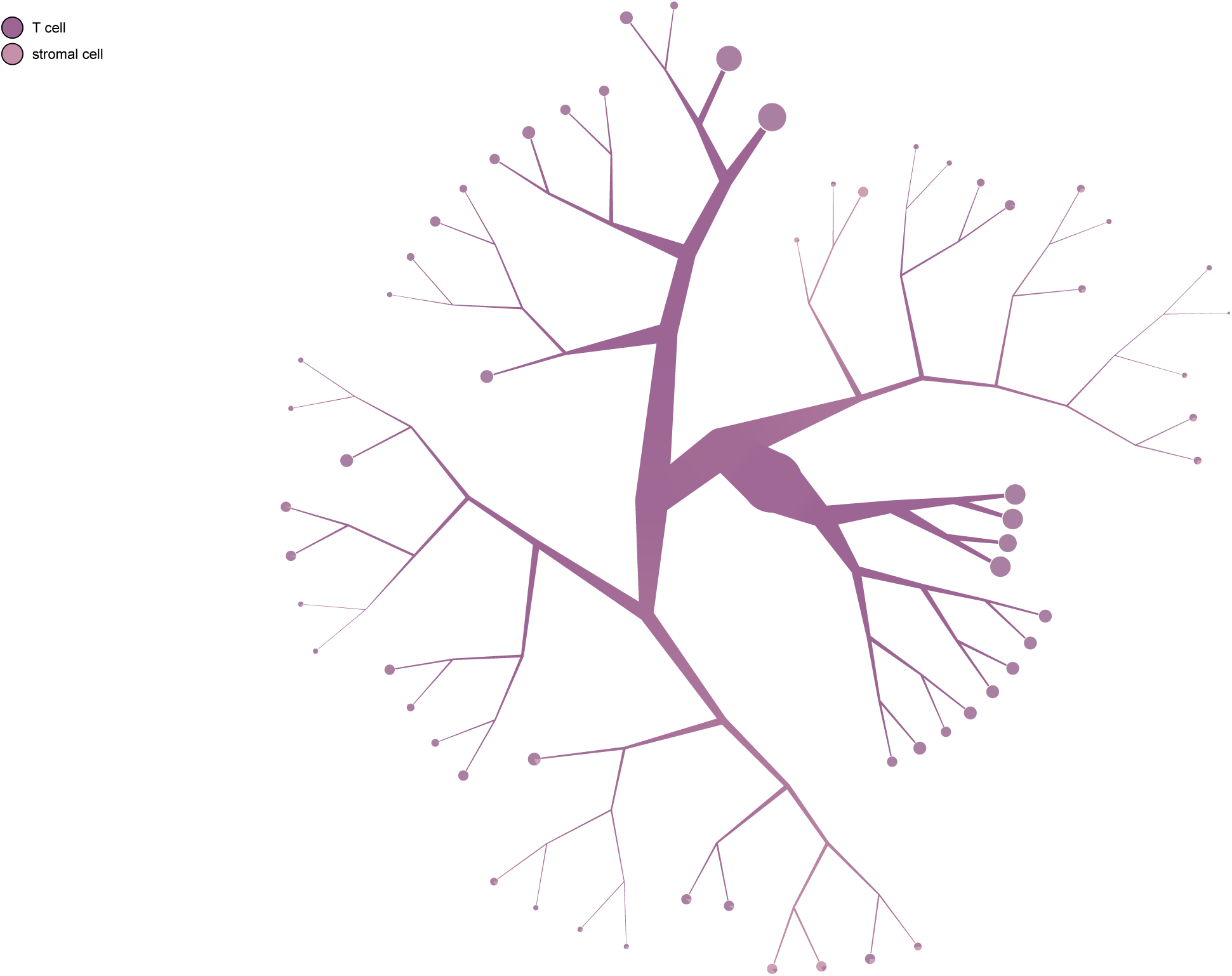
`TooManyCells` tree with cell type overlay in thymus.

**Figure S17:**
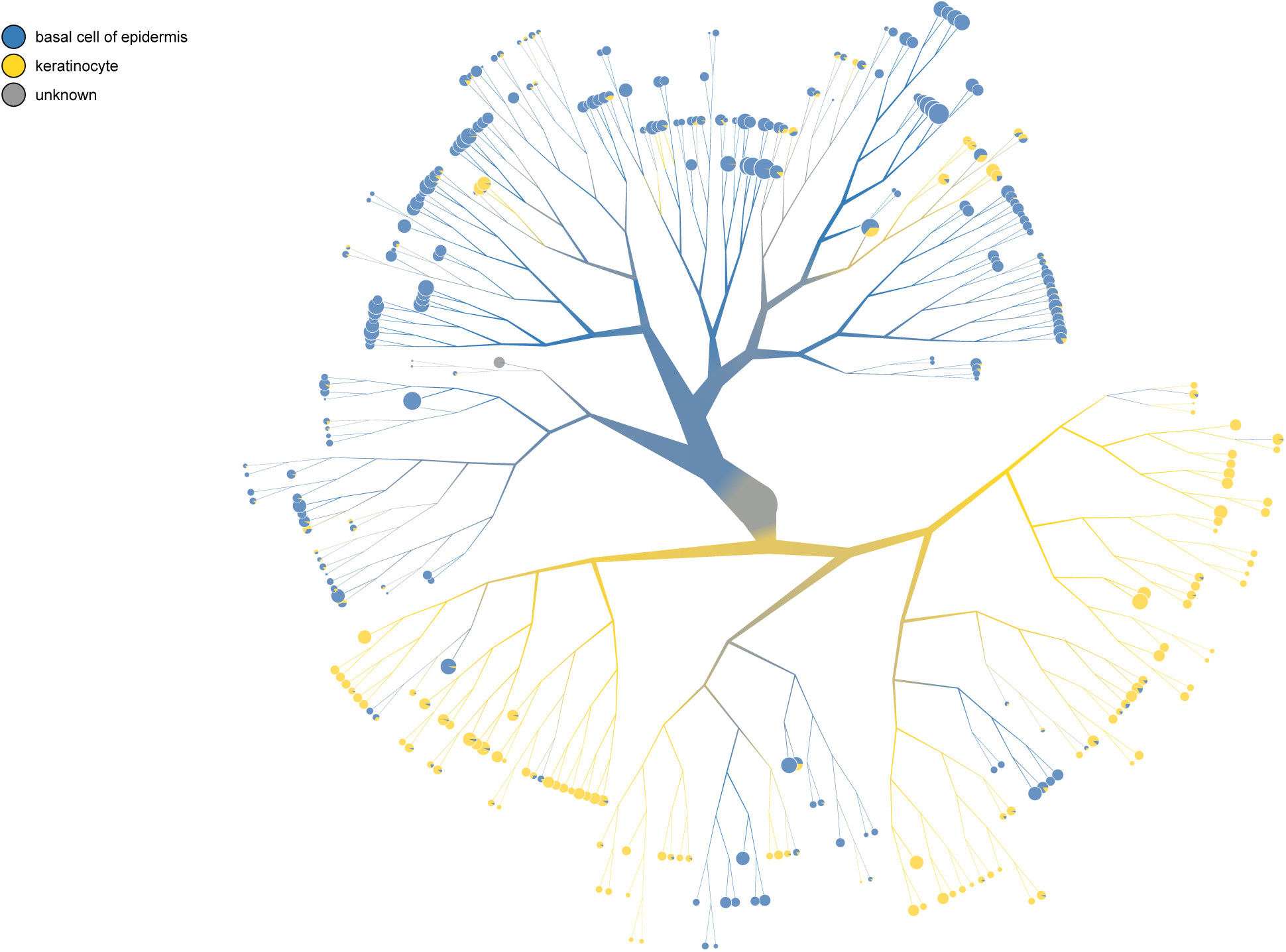
`TooManyCells` tree with cell type overlay in tongue.

**Figure S18:**
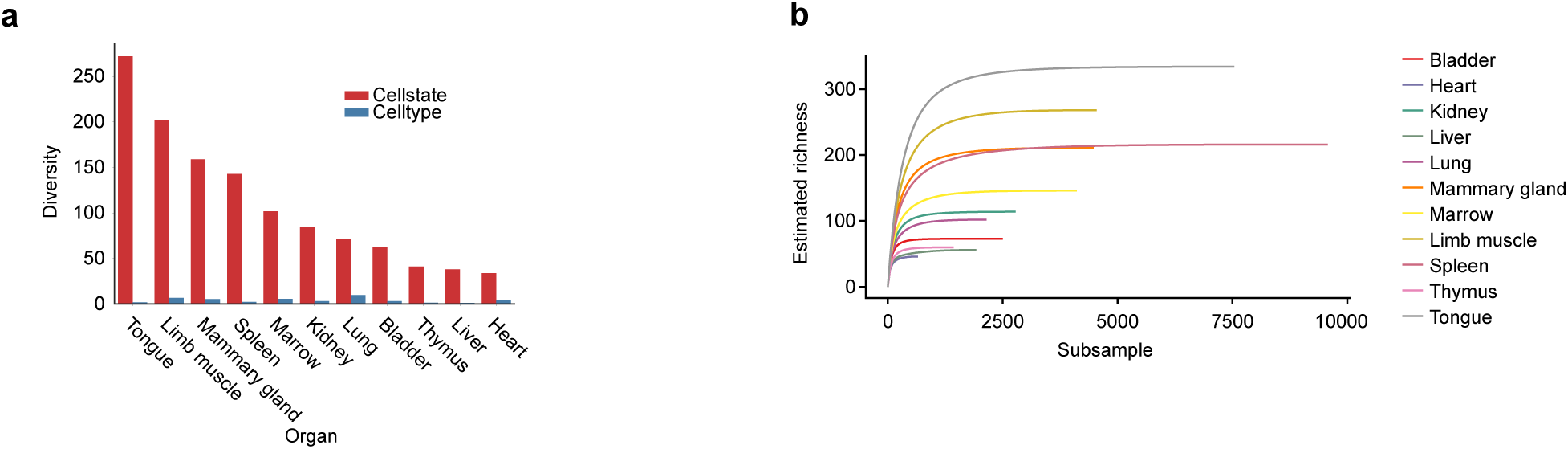
Diversity of organs from the `TooManyCells` tree leaves and by cell celltype. (a) The diversity of each organ by most fine-grain cell state (red, `TooManyCells` leaves) and by previous annotations (blue). (b) Rarefaction curves of the most fine-grain cell state for each organ.

**Figure S19:**
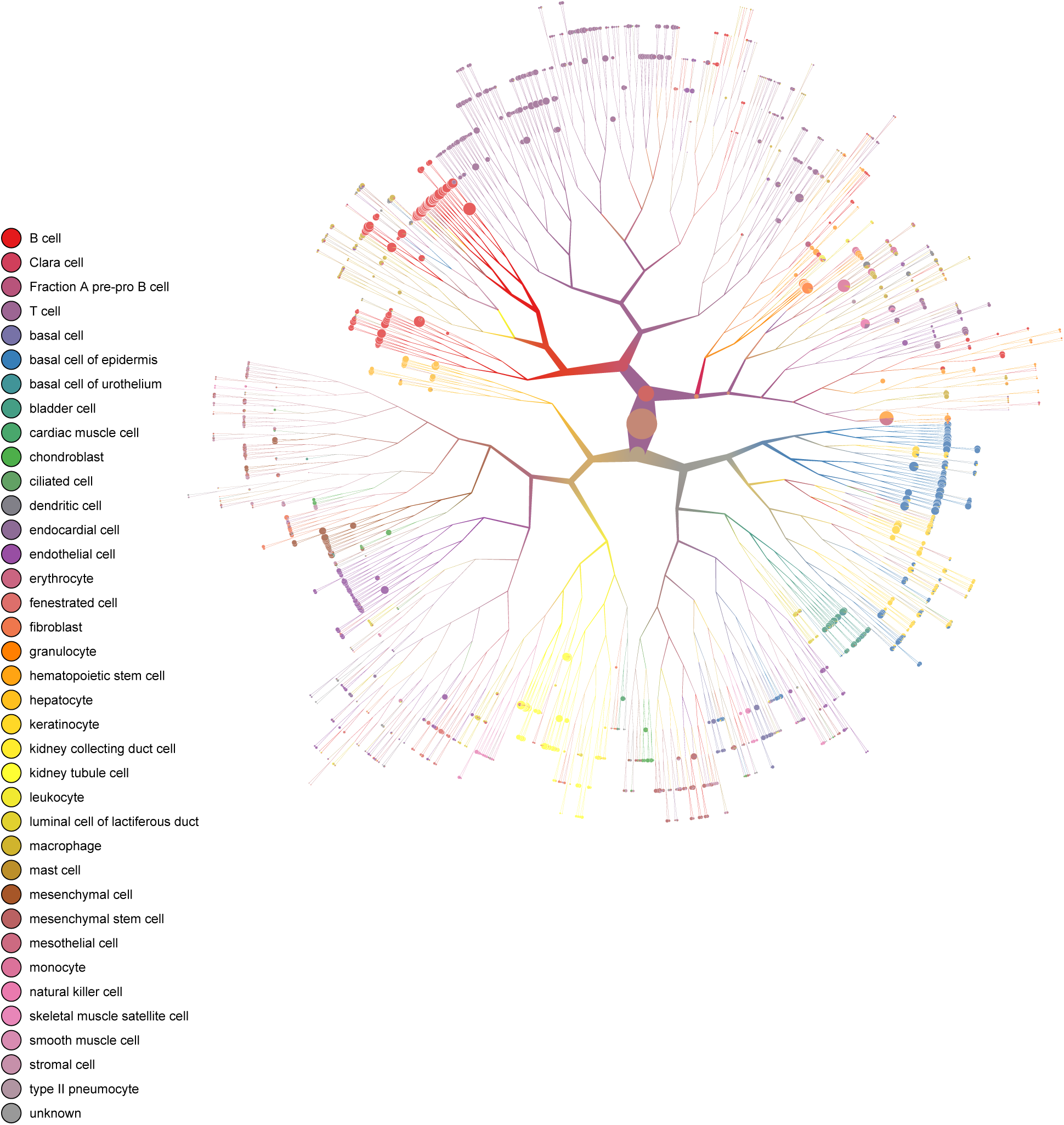
`TooManyCells` tree with cell type overlay present within 11 organs and tissues.

**Figure S20:**
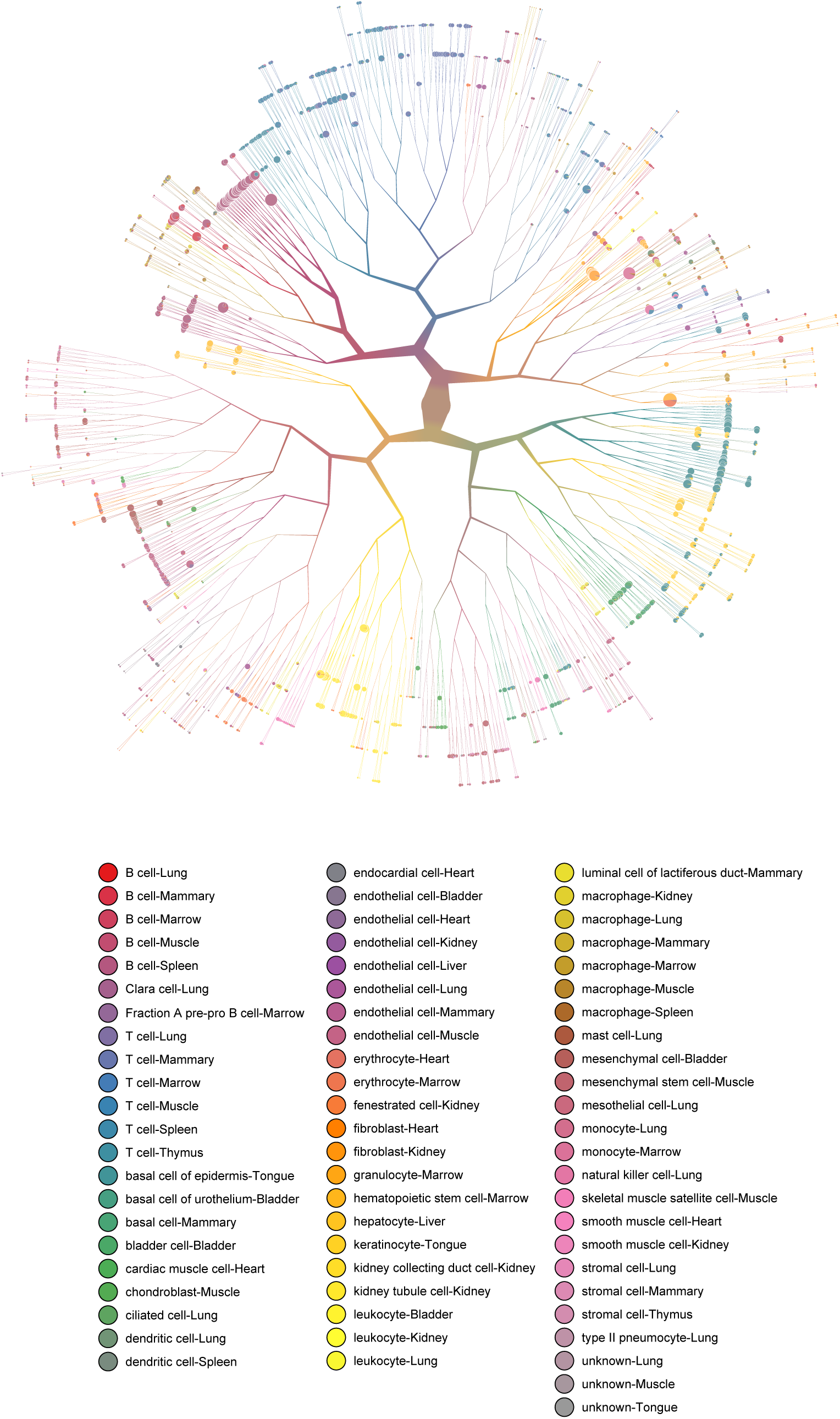
`TooManyCells` tree with cell type and each tissue overlay present within 11 organs and tissues.

**Figure S21:**
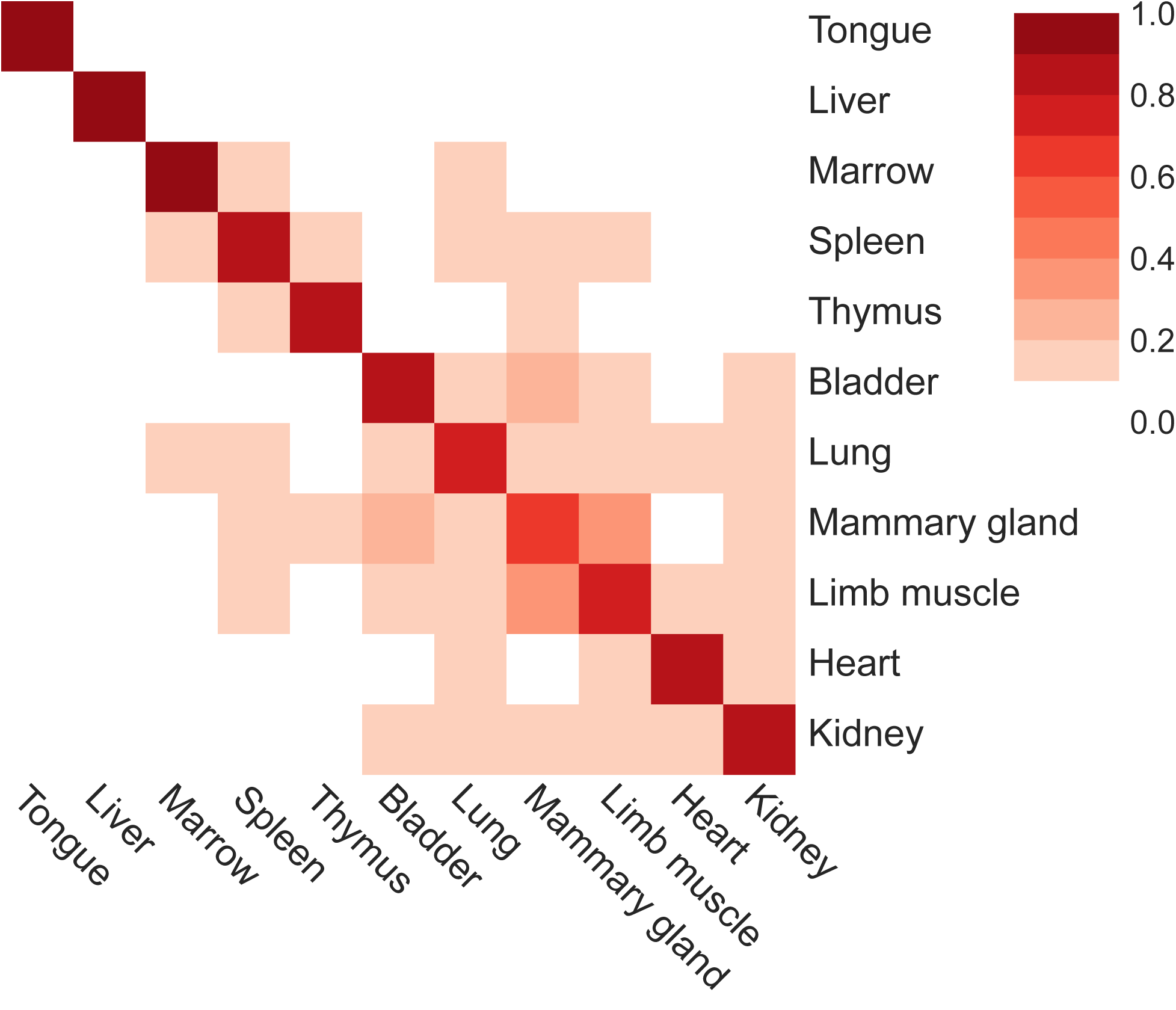
Clumpiness of 11 organs within the `TooManyCells` tree.

**Figure S22:**
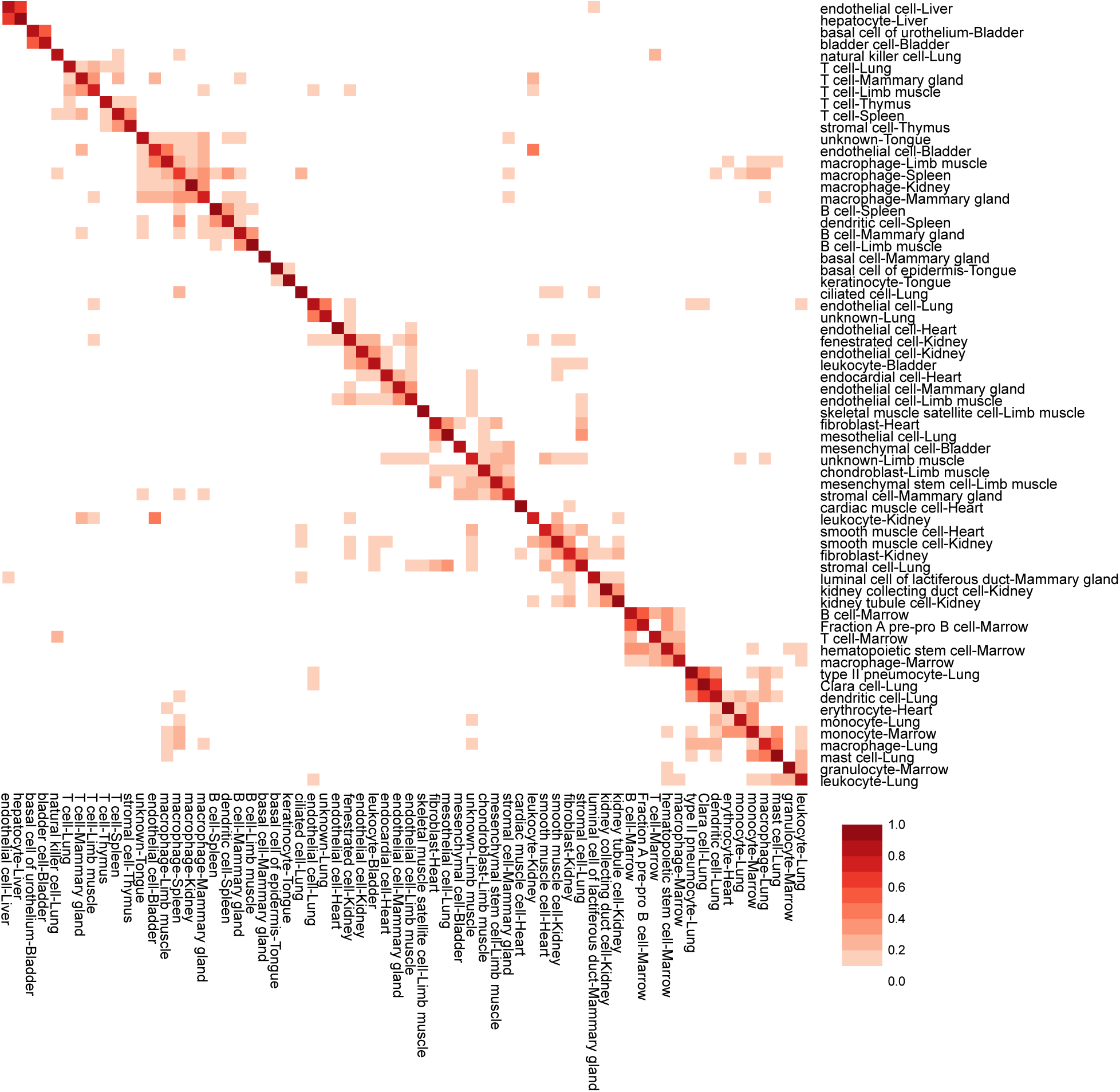
Clumpiness of cell types with respective organs from 11 organs within the `TooManyCells` tree.

**Figure S23:**
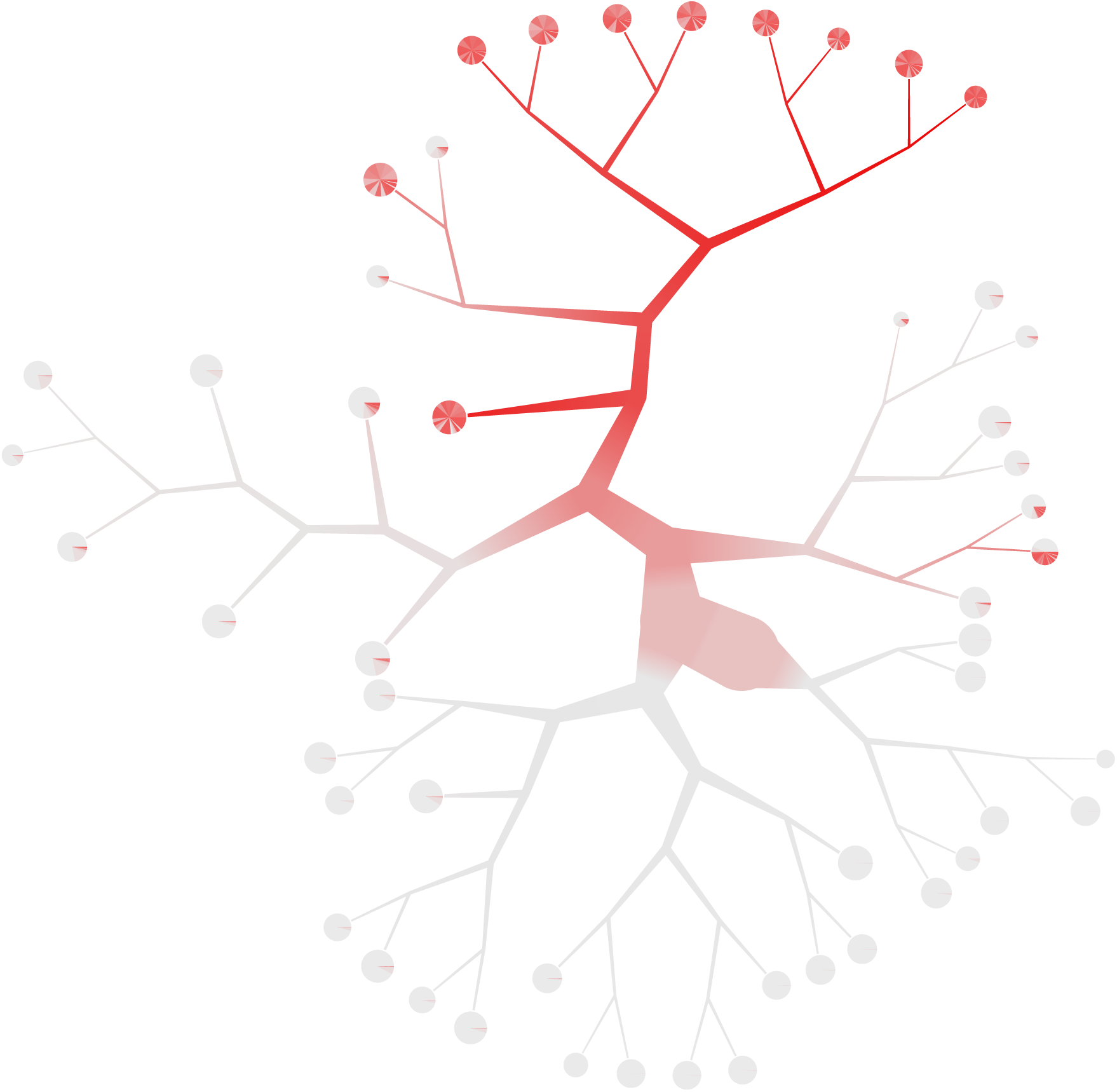
`TooManyCells` tree with cutting criterion `--smart-cutoff 4` and *Cd79a* expression as a marker for B cells from low (white) to high (red).

